# Epithelial micro-invasion by *Streptococcus pneumoniae* induces epithelial-derived innate immunity during colonisation at the human mucosal surface

**DOI:** 10.1101/281576

**Authors:** Caroline M Weight, Cristina Venturini, Sherin Pojar, Simon P. Jochems, Jesús Reiné, Elissavet Nikolaou, Carla Solórzano, Mahdad Noursadeghi, Jeremy S Brown, Daniela M. Ferreira, Robert S Heyderman

## Abstract

Control of *Streptococcus pneumoniae* colonisation at human mucosal surfaces is critical to reducing the burden of pneumonia and invasive pneumococcal disease, interrupting transmission, and achieving herd protection. Using an Experimental Human Pneumococcal Carriage Model (EHPC), we show that *S. pneumoniae* colonisation is associated with epithelial surface adherence, micro-colony formation and invasion, without overt disease. Interactions between different strains and the epithelium *in vitro* shaped the host transcriptomic response. Using epithelial modules from a human epithelial cell model that recapitulates our *in vivo* findings, comprising of innate signalling/ regulatory pathways, inflammatory mediators, cellular metabolism and stress response genes, we find that inflammation in the EHPC model is most prominent around the time of bacterial clearance. These results show that rather than being confined to the epithelial surface and the overlying mucus layer, the pneumococcus undergoes micro-invasion of the epithelium that enhances the inflammatory/ innate immune response associated with clearance.

## INTRODUCTION

Colonisation of upper respiratory tract (URT) mucosa by a range of pathogenic bacteria is a necessary precursor to both disease and onward transmission. Although *Streptococcus pneumoniae* is a common coloniser of the human nasopharynx, it is estimated to be responsible for >500,000 deaths due to pneumonia, meningitis and sepsis in children under five years of age worldwide^1^.

In Europe and North America, there has been a dramatic effect of pneumococcal conjugate vaccine (PCV) on vaccine serotype (VT) invasive disease, carriage and transmission^2^. However, the emergence of non-VT pneumococcal disease worldwide and the more modest impact of PCV on colonisation in high transmission settings, threaten this success^3^. Control of pneumococcal URT colonisation in humans is not fully understood^4^, and so defining the mechanistic basis for host control of pneumococcal colonisation at the mucosal surface is therefore crucial for the further optimisation of therapeutic interventions which target carriage and transmission. The process of transmission is not fully understood, but mucosal inflammation, potentially enhanced by co-infection with viruses such as Influenza A, has been proposed to mediate bacterial shedding from the nasopharynx^5^.

Naturally acquired immunity to *S. pneumoniae* proteins are primarily mediated by mucosal T cells and anti-protein antibodies, controlled by Treg^6-8^. The role of anti-capsule polysaccharide antibodies in naturally acquired immune control remains unresolved^9^. URT epithelium is central to this immunity^10^, binding and transporting antibodies, sensing bacteria via a range of surface and intracellular pathogen-associated molecular patterns (PAMPs) receptors, and rapidly transducing signals to recruit innate and inflammatory immune mechanisms^11,12^.

Murine models suggest that adherence of *S. pneumoniae* to the mucosal epithelium may be followed by paracellular transmigration and tight junction modulation^13,14^. In contrast, studies of immortalised epithelial cell monolayers implicate endocytosis of *S. pneumoniae*, mediated though pneumococcal protein C–polymeric immunoglobulin receptor interactions^15,16^. The relative importance of epithelial endocytosis and paracellular migration in colonisation and invasion remains uncertain^16^, but could influence epithelial sensing of this otherwise extracellular pathogen^14^. For example this may include Nod1 signalling via peptidoglycan^17^ and TLR4 signalling via pneumolysin, a pore-forming toxin that induces inflammation and mediates both clearance and transmission in an infant mouse model^18-20^.

We have explored the hypothesis that pneumococcal-epithelial engagement dictates the inflammatory/ innate immune response and therefore the outcome of colonisation. Much of what we understand of the control of pneumococcal colonisation is derived from epidemiological studies and murine carriage models. The Experimental Human Pneumococcal Carriage (EHPC) model provides a well-controlled, reproducible tool to characterise the cellular and molecular mechanisms that underlie pneumococcal colonisation in humans^21^. The model has revealed that pneumococcal carriage results in neutrophil degranulation and monocyte recruitment to the nasopharynx^22^, protecting against re-challenge with the same strain for up to one year^21,23^.

We have used human epithelial cell lines and different strains of *S. pneumoniae* to further probe these mechanisms and derive an epithelial transcriptome module to interrogate the host response in the EHPC model. We show that pneumococcal colonisation in humans is characterised by micro-colony formation, junctional protein association and migration across the epithelial barrier without disease, which we have termed micro-invasion. This pattern of bacterial-host cell association shapes epithelial sensing of *S. pneumoniae*, which is partially pneumolysin dependent. Together, our data suggest that pneumococcal engagement with the epithelium in the early phases of colonisation may occur without eliciting a marked host response, but as colonisation becomes more established, epithelial sensing of *S. pneumoniae* enhances innate immunity/ inflammation which we propose promotes clearance.

## RESULTS

### Pneumococcal colonisation of the human nasopharynx is associated with epithelial adhesion, micro-colony formation and micro-invasion

We have used an EHPC model^21^ to capture the epithelial events central to pneumococcal colonisation, obtaining human mucosal samples by curette biopsy. We defined a carrier as an individual colonised by 6B *S. pneumoniae* as detected by culture of nasal wash at any day following inoculation. Clearance was defined as when *S. pneumoniae* colonisation became negative by culture of nasal wash.

Colonisation was detected in 9/13 healthy volunteers by culture, 12/13 by microscopy (using 6B capsule-specific antibody labelling) and 9/11 by LytA PCR (Table 1). Carriage was detected by culture/ PCR at day 2, whereas maximum detection by microscopy was evident at day 6. This is not explained by colonisation density and therefore the sensitivity of culture/ PCR. This could be explained by greater association with the epithelium (detected by all techniques) rather than the overlying mucus (detected by culture/ PCR but not microscopy) over time. Clearance generally occurred between day 9 and day 27 (Table 1 and Figure 1b).

**Table 1.** *Streptococcus pneumoniae* association with the nasal epithelium in the EHPC model detected by culture, confocal microscopy and LytA PCR. Nasal washes and nasal curette biopsies were collected from carriage positive and carriage negative volunteers over time. Standard methods for measuring bacterial density by culture (CFU) and LytA PCR were compared against counts visualised by confocal microscopy for pneumococcal association with nasal cells over time. The data was derived from 13 volunteers. + (1-10 pneumococci); ++ (11-50 pneumococci); +++ (51-100 pneumococci); ++++ (>100 pneumococci); n/a = reading not taken.

| ID | Pneumococcal density | Sample day |  |  |  |  | Pneumococcal association |  |  |
| --- | --- | --- | --- | --- | --- | --- | --- | --- | --- |
|  |  | Pre | D2 | D6 | D9 | D27 | Surface | Intracellular | Lateral |
| 1 | Culture (CFU/ml) | 0 | 373 | 520 | 10 | 2 |  |  |  |
|  | Microscopy (counts) | 0 | + | + | 0 | + | + | 0 | 0 |
|  | LytA PCR | NEG | POS | POS | n/a | POS |  |  |  |
| 2 | Culture (CFU/ml) | 0 | 0 | 0 | 0 | 0 |  |  |  |
|  | Microscopy (counts) | + | + | 0 | 0 | 0 | ++ | + | 0 |
|  | LytA PCR | NEG | NEG | NEG | n/a | NEG |  |  |  |
| 3 | Culture (CFU/ml) | 0 | 0 | 0 | 0 | 0 |  |  |  |
|  | Microscopy (counts) | 0 | + | 0 | 0 | 0 | + | 0 | 0 |
|  | LytA PCR | NEG | POS | NEG | n/a | NEG |  |  |  |
| 4 | Culture (CFU/ml) | 0 | 3 | 9 | 0 | 0 |  |  |  |
|  | Microscopy (counts) | 0 | 0 | 0 | + | 0 | + | 0 | 0 |
|  | LytA PCR | NEG | POS | POS | n/a | NEG |  |  |  |
| 5 | Culture (CFU/ml) | 0 | 7x10<sup>3</sup> | 223 | 22 | 0 |  |  |  |
|  | Microscopy (counts) | 0 | + | + | + | 0 | + | + | 0 |
|  | LytA PCR | n/a | n/a | n/a | n/a | n/a |  |  |  |
| 6 | Culture (CFU/ml) | 0 | 1.9x10<sup>4</sup> | 5.7x10<sup>3</sup> | 3.7x10<sup>4</sup> | 11 |  |  |  |
|  | Microscopy (counts) | 0 | ++ | ++++ | ++ | 0 | +++ | ++ | + |
|  | LytA PCR | NEG | POS | POS | n/a | POS |  |  |  |
| 7 | Culture (CFU/ml) | 0 | 822 | 1.1x10<sup>3</sup> | 160 | 0 |  |  |  |
|  | Microscopy (counts) | 0 | + | +++ | ++ | 0 | ++ | + | + |
|  | LytA PCR | NEG | POS | POS | n/a | NEG |  |  |  |
| 8 | Culture (CFU/ml) | 0 | 0 | 0 | 0 | 0 |  |  |  |
|  | Microscopy (counts) | 0 | 0 | 0 | 0 | 0 | 0 | 0 | 0 |
|  | LytA PCR | n/a | n/a | n/a | n/a | n/a |  |  |  |
| 9 | Culture (CFU/ml) | 0 | 221 | 695 | 4 | <1 |  |  |  |
|  | Microscopy (counts) | n/a | + | +++ | + | n/a | +++ | + | + |
|  | LytA PCR | NEG | POS | POS | n/a | NEG |  |  |  |
| 10 | Culture (CFU/ml) | 0 | 1x10<sup>3</sup> | 0 | 0 | 0 |  |  |  |
|  | Microscopy (counts) | n/a | 0 | + | 0 | n/a | + | 0 | 0 |
|  | LytA PCR | NEG | POS | POS | n/a | POS |  |  |  |
| 11 | Culture (CFU/ml) | 0 | 1.8x10<sup>6</sup> | 1.8x10<sup>6</sup> | 1.9x10<sup>4</sup> | <1 |  |  |  |
|  | Microscopy (counts) | n/a | ++++ | ++++ | + | n/a | +++ | + | + |
|  | LytA PCR | NEG | POS | POS | n/a | NEG |  |  |  |
| 12 | Culture (CFU/ml) | 0 | 0 | 0 | 0 | 0 |  |  |  |
|  | Microscopy (counts) | n/a | ++ | 0 | 0 | n/a | ++ | + | + |
|  | LytA PCR | NEG | NEG | NEG | n/a | NEG |  |  |  |
| 13 | Culture (CFU/ml) | 0 | 23 | 2 | 0 | 0 |  |  |  |
|  | Microscopy (counts) | n/a | + | 0 | 0 | n/a | + | 0 | 0 |
|  | LytA PCR | POS | POS | POS | n/a | POS |  |  |  |

**Figure 1.**
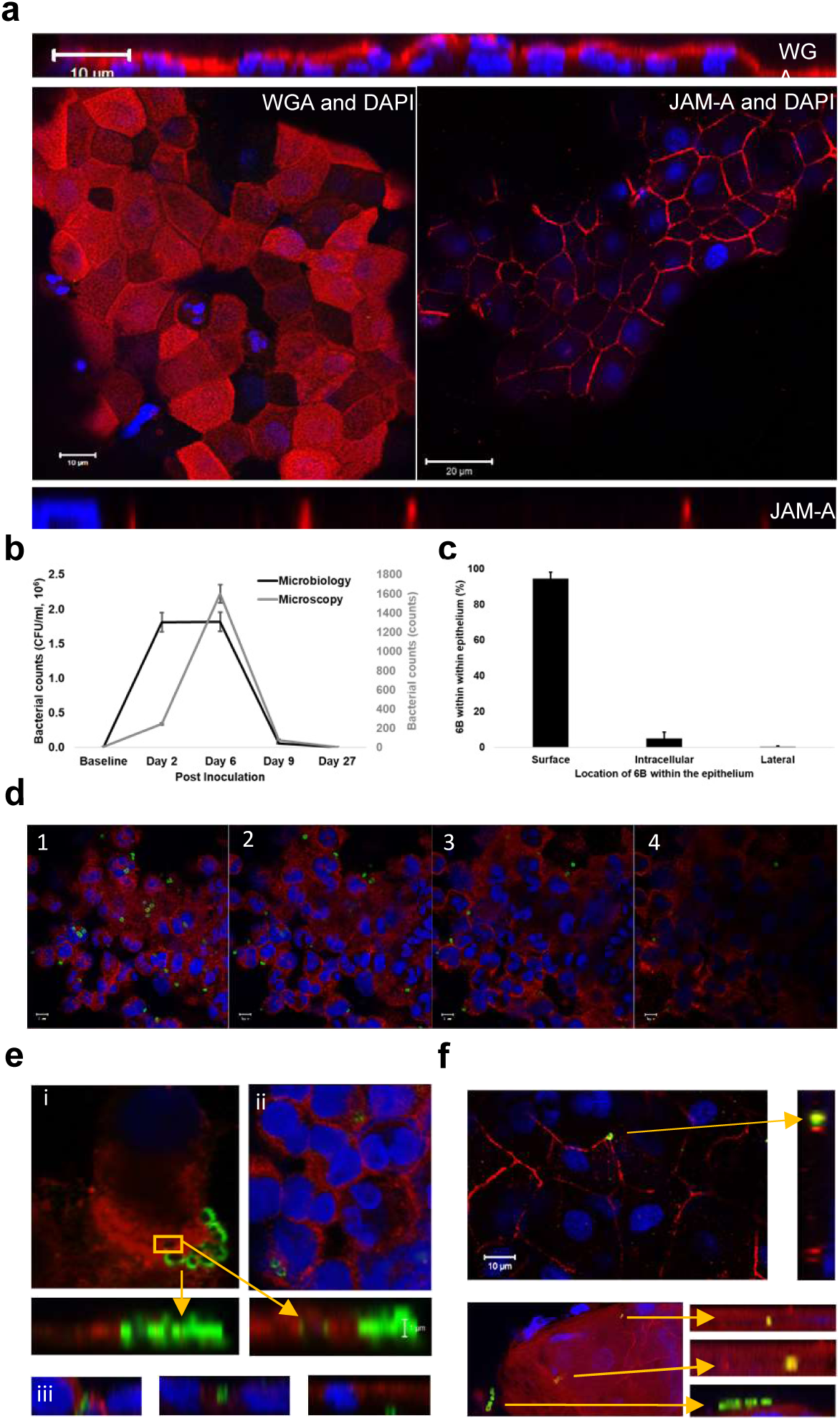
*Streptococcus pneumoniae* colonisation of the human nasal epithelium from the EHPC model is associated with adhesion, micro-colony formation and micro-invasion in response to *S. pneumoniae* 6B colonisation. (a) Representative fields from the EHPC model nasal curette biopsies show intact epithelium can be obtained from the samples (as visualised by XY planes) and they retain their size and shape (as visualised by XZ planes). Cells were stained with Wheat Germ Agglutinin (WGA) or Junctional Adhesion Molecule A (JAM-A, red) and nuclei (blue). (b) The pattern of pneumococcal density detected by culture and microscopy. (c) The proportion of bacteria located on the cell surface, intracellularly, or paracellularly visualised by confocal microscopy, quantified from 8 volunteers collected over time from microscopy counts. (d) Cells were stained for surface carbohydrates using WGA (red), and the bacteria were marked with specific serotype antiserum (green). XY images of 1μm slices through a layer of cells from top (stack 1) to bottom (stack 4), with bacteria associated. (e) Examples of cells collected on day 6 post-inoculation stained with WGA (red) showing; (ei) mMicro-colony formation on the surface of the epithelium, (eii) bacterial internalization, (eiii) migration through epithelial uptake internally or between cells. (f) Co-association between *S. pneumoniae* (green) and JAM-A (red). Nuclei (blue). Bacterial uptake appeared co-associated with JAM-A (yellow).

Curette biopsy samples visualised by confocal microscopy using Wheat Germ Agglutinin (WGA) as a marker for surface carbohydrates and Junctional Adhesion Molecule A (JAM-A) as a marker of tight junctions, yielded intact sheets of epithelial cells (Supplementary Figure 1bi, 1bii) associated with polymorphonuclear leukocytes, characterised morphologically (Figure 1a and Supplementary Figure 1a). Amongst carriers, pneumococci were found on the epithelial surface (Table 1; Figure 1c, Figure 1d, Supplementary Figure 1biii) which over time (typically by 6 days), consisted of two or more pneumococci at the same location on the epithelial surface, which we have termed micro-colonies (Figure 1e, Supplementary Figure 1biv). There was evidence of what we have termed pneumococcal “micro-invasion” of the epithelial monolayer consisting of either internalisation of bacteria and/ or transmigration across the epithelial barrier (Figure 1e-f). Pneumococci were also observed in phagocytic cells (Supplementary Figure 1a). Colonisation was frequently characterised by co-association between *S. pneumoniae* and JAM-A (Figure 1f). This intimate association between inoculated pneumococci and the mucosa during asymptomatic pneumococcal carriage in humans is suggestive of an active engagement process which may influence the outcome of colonisation.

Further visualization of this bacteria-host interaction by transmission electron microscopy has proven problematic, but we have been able to demonstrate the integrity of the nasal epithelial curette biopsy samples (Supplementary Figures 1bi and 1bii). We provide evidence of diplococci on the epithelial cell surface and chains or micro-colonies of diplococci that may have been dislodged from the epithelial surface (Supplementary Figures 1biii and 1biv).

### Epithelial surface marker expression *in vivo* is not altered in response to colonisation with *S. pneumoniae*

To determine whether pneumococcal colonisation leads to epithelial activation, we analysed nasal curette biopsy cells for surface expression of molecules expressed in airway epithelium which have previously been shown to be upregulated in response to common bacterial stimuli and are involved in immune cell recruitment. These were EpCAM (for epithelial cell identification), IL-22Ra1, HLADR, CD40, CD54 and CD107a (Supplementary Figure 2a-d). We did not detect a significant change in the relative expression of IL-22Ra1 (protects the epithelial barrier, promotes anti-microbial product secretion during infection, modulating pneumococcal carriage and clearance^24^), HLADR (mediates T cell-antigen recognition and is a marker of epithelial activation^25^), CD40 (co-stimulatory protein which binds CD154^26^ (Supplementary Figure 2e-g)), or CD54 (a leukocyte adhesion molecule which is also upregulated by CD40^26,27^, initiating neutrophil migration and recruitment^28^, Figure 2a). However, although numbers of epithelial cells expressing CD107a did not change over time (Figure 2b, left), we observed an increase in CD107a^high^ expression at day 2 post inoculation in carriage positive volunteers vs. carriage negative volunteers (Figure 2b, right). CD107a in the epithelium forms the membrane glycoprotein of late endosomes and phagolysosomes^16^ and has previously been implicated in pneumococcal endocytosis^16^. These data highlight the ability of the pneumococcus to colonise the nasopharynx without marked inflammation and potentially implicate CD107a in the process.

**Figure 2.**
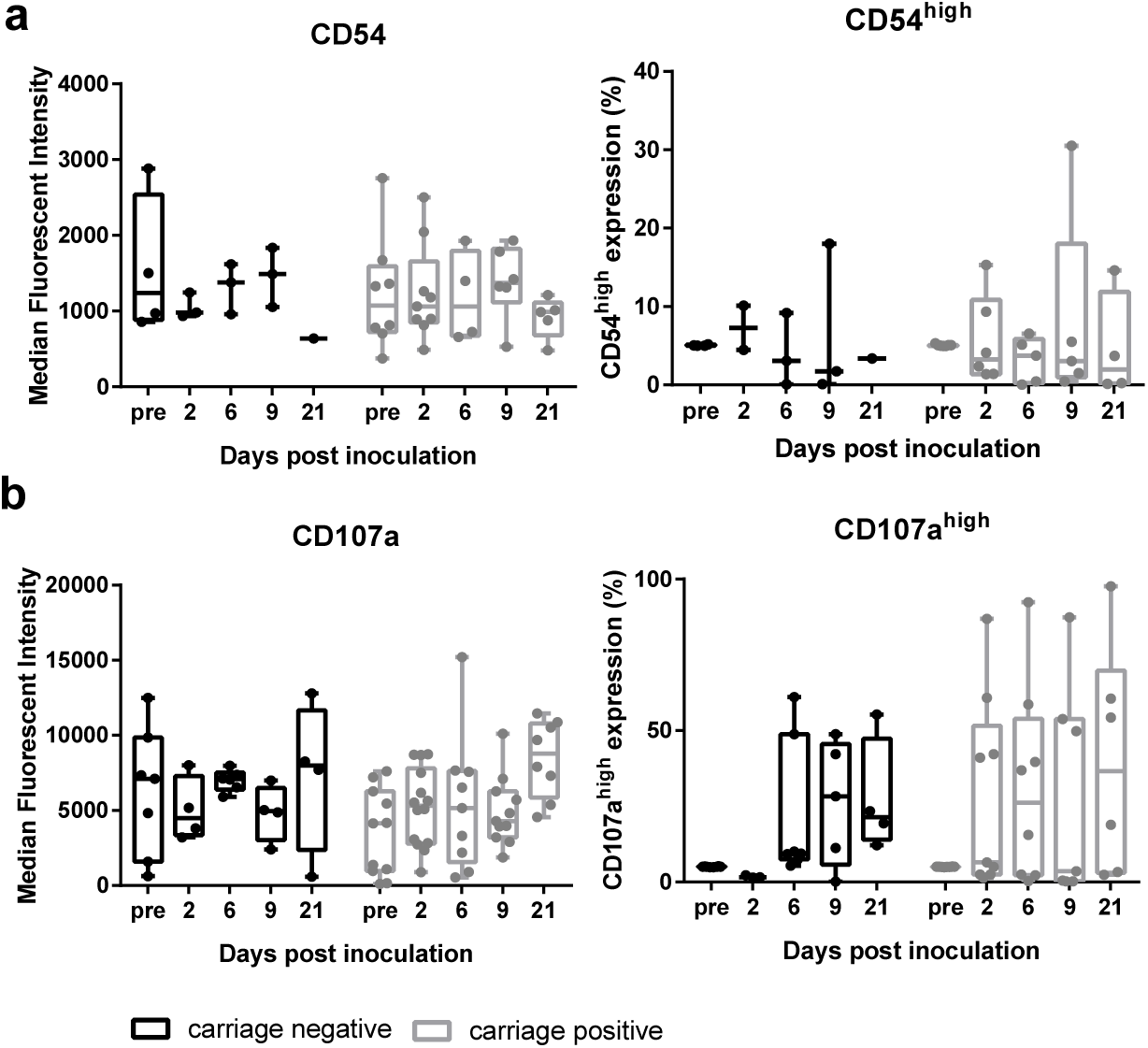
Epithelial surface marker expression is not altered in response *to S. pneumoniae* 6B colonisation *in vivo*. Epithelial cells from nasal curette biopsies were analysed by flow cytometry for (a) CD54 and (b) CD107a. Results were expressed as median fluorescence intensity (left) and high cell surface expression (≥95% of the baseline expression, right) representing at least two volunteers per time point who were carriage negative (black) and carriage positive (grey). Comparisons between carriage negative and carriage positive, on each day of sampling, was analysed by 1-way ANOVA (a – parametric, b – non-parametric). P = 0.84 CD54 MFI; p = 0.98 CD54 %; p = 0.09 CD107a MFI; p = 0.10 CD107a %.

### *S. pneumoniae* infection of nasopharyngeal epithelial cells *in vitro* is associated with pneumococcal adhesion, micro-colony formation and micro-invasion

To further understand epithelial responses to *S. pneumoniae* observed in the EHPC model, we undertook infection experiments with Detroit 562 epithelial cells. We compared the 6B response with two other representative clinical isolates; a 23F strain, TIGR4 (the original sequenced strain^29^) and a TIGR4 pneumolysin deletion mutant strain (dPly).

TIGR4 association with Detroit 562 cells was over 100-fold higher for adhesion and over 70-fold higher for invasion after three hours infection, compared to 6B or 23F strains (Figure 3a-b). Using capsule-specific antibody stain for confocal and electron microscopy, pneumococcal adhesion to the epithelial surface at 3 hours was found to be associated with micro-colony formation, pneumococcal chain formation and internalisation of this conventionally extracellular pathogen^14^ (Figure 3d-e, Figure i-j, Supplementary Figure 3a). Internalisation was associated with the formation of epithelial membrane folds (Figure 3j left). Intracellular pneumococci were contained within intracellular vesicles suggestive of endocytosis, which was most marked with the TIGR4 and dPLY-TIGR4 strains (Figure 3b, 3j)^16^. Intracellular pneumococci were seen within vacuoles (Figure 3j middle), coated with JAM-A (a tight junction protein that regulates barrier function and cell polarity, Figure 3f, Supplementary Figure 3a) and associated with the adherens junction protein β catenin (that is involved in epithelial adhesion), but not Claudin 4 (a tight junction protein that regulates paracellular permeability, Supplementary Figure 3a). Pneumococci were located laterally, transiting between cell junctions (Figure 3g, 3j right), at the level of the nuclei and, below the basal membrane (Figure 3h). Transmigration across the epithelium (endocytosis or paracellular movement), as measured by the culture of pneumococci from the basal chamber of transwell inserts, was most marked with 23F (Figure 3c, five-fold higher compared to TIGR4, after three hours infection). A similar pattern of epithelial interaction was also seen at one hour suggesting these observations are not simply explained by differential growth.

**Figure 3.**
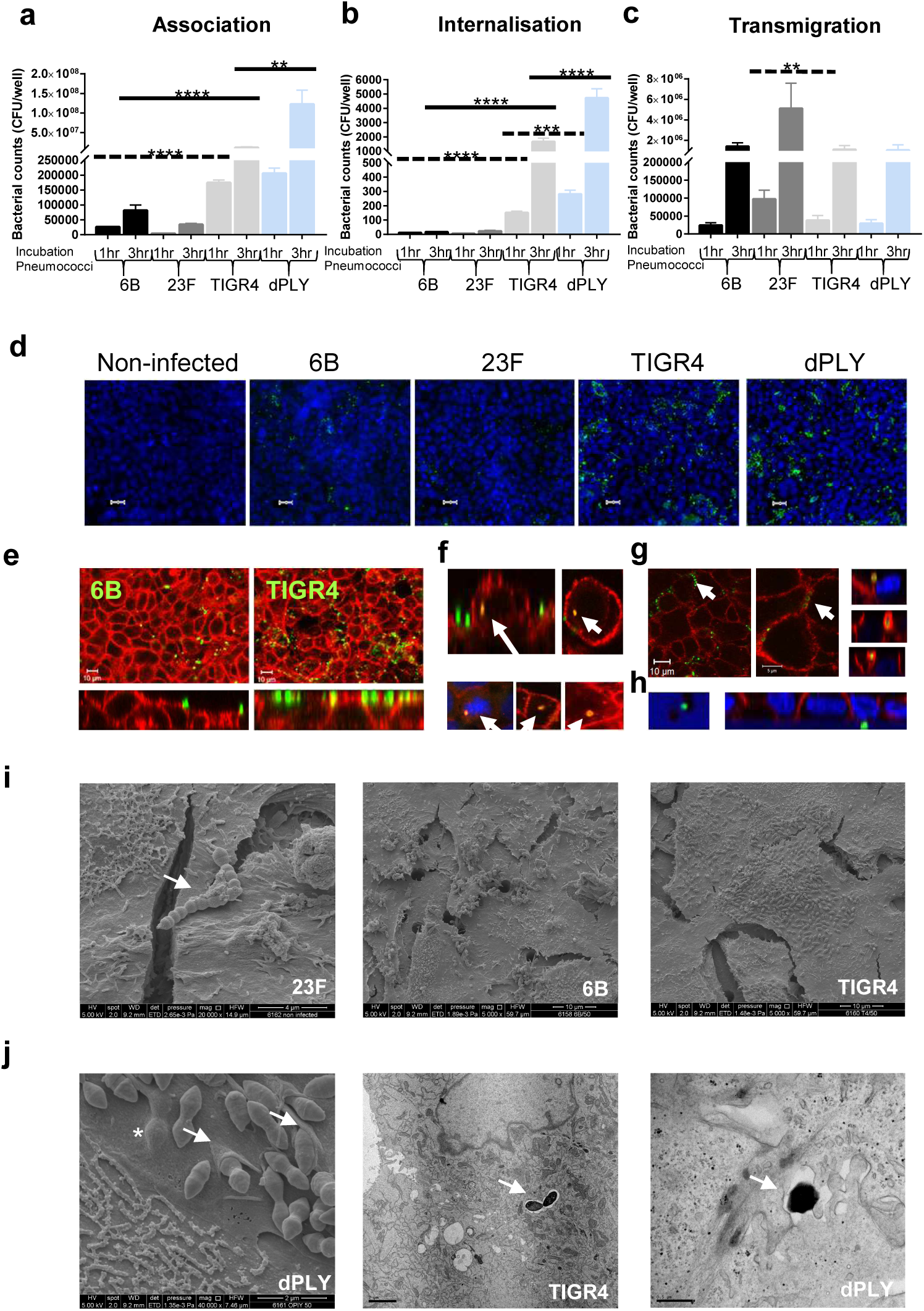
*Streptococcus pneumoniae* infection of nasopharyngeal epithelium *in vitro* is associated with adhesion, micro-colony formation, micro-invasion, transmigration and is partially influenced by pneumolysin. (a) Pneumococcal association and (b) internalisation: Detroit 562 cell monolayers were stimulated with pneumococci for 1 or 3 hours and the quantity of associated bacteria was determined by culture (CFU); n = 6. ****P = <0.0001, ***P = 0.0001: – (a) 1hr (dashed lines, ANOVA); 3hr (solid lines, Kruskall-Wallis); (b) (Kruskall-Wallis). (c) Pneumococcal transmigration: Cells on transwell inserts were stimulated with pneumococci for 3 hours and bacterial density in the basal chamber was determined (**P = 0.0092, Kruskall-Wallis). N = 5. (d) Representative pneumococcal-density (green) images of cells (blue nuclei). Scale bar = 20μm. n = 5. (e – h) Representative images of pneumococcal localization (green) from cells (red, JAM-A), illustrating: (e) differences in adherence by 6B and TIGR4; (f) internalised bacteria (top 6B; bottom TIGR4) co-localised with JAM-A with associated intracellular vesicle-like bodies (yellow); (g) lateral localisation of pneumococci (XY images, TIGR4) with possible paracellular movement co-associated with JAM-A (XZ images, 6B); (h) basal localisation of bacteria (23F) at the level of nuclei and insert pores N = 5 with replicates. (i) Scanning EM images of Detroit 562 cells infected with *S. pneumoniae* which appear as diplococci. Left – pneumococcal chain formation (strain 23F, arrow); middle and right – micro-colony formation by strains 6B and TIGR4 on the epithelial surface. (i) Micro-invasion of dPLY-TIGR4 shown by EM: left – scanning EM showing epithelial membrane folding (arrows) and pneumococci below the cell membrane surface (*); middle – internalisation of TIGR4 pneumococci encased within a vacuole (arrow); and right – transmission EM showing transmigration of dPLY-TIGR4 pneumococci between epithelial cells (arrow).

Epithelial infection with the dPly-TIGR4 mutant showed a significant increase in adherence (Figure 3a, ten-fold higher after three hours infection), internalisation (Figure 3b, three-fold higher after three hours infection, Figure 3j), but no significant differences in transmigration (Figure 3c) compared to the wild-type strain, implicating pneumolysin in the modulation of bacterial interactions with the epithelium.

Similar patterns of adherence and invasion were observed in infection experiments with human Calu3 and A549 epithelial cells demonstrating that these findings are cell-line independent (Supplementary Figure 4). Furthermore, primary epithelial cells differentiated on an air-liquid interface for 30 days and co-cultured with either 6B or 23F *S. pneumoniae* also revealed pneumococcal micro-colony formation, micro-invasion and epithelial junctional protein association (Supplementary Figure 1c).

**Figure 4.**
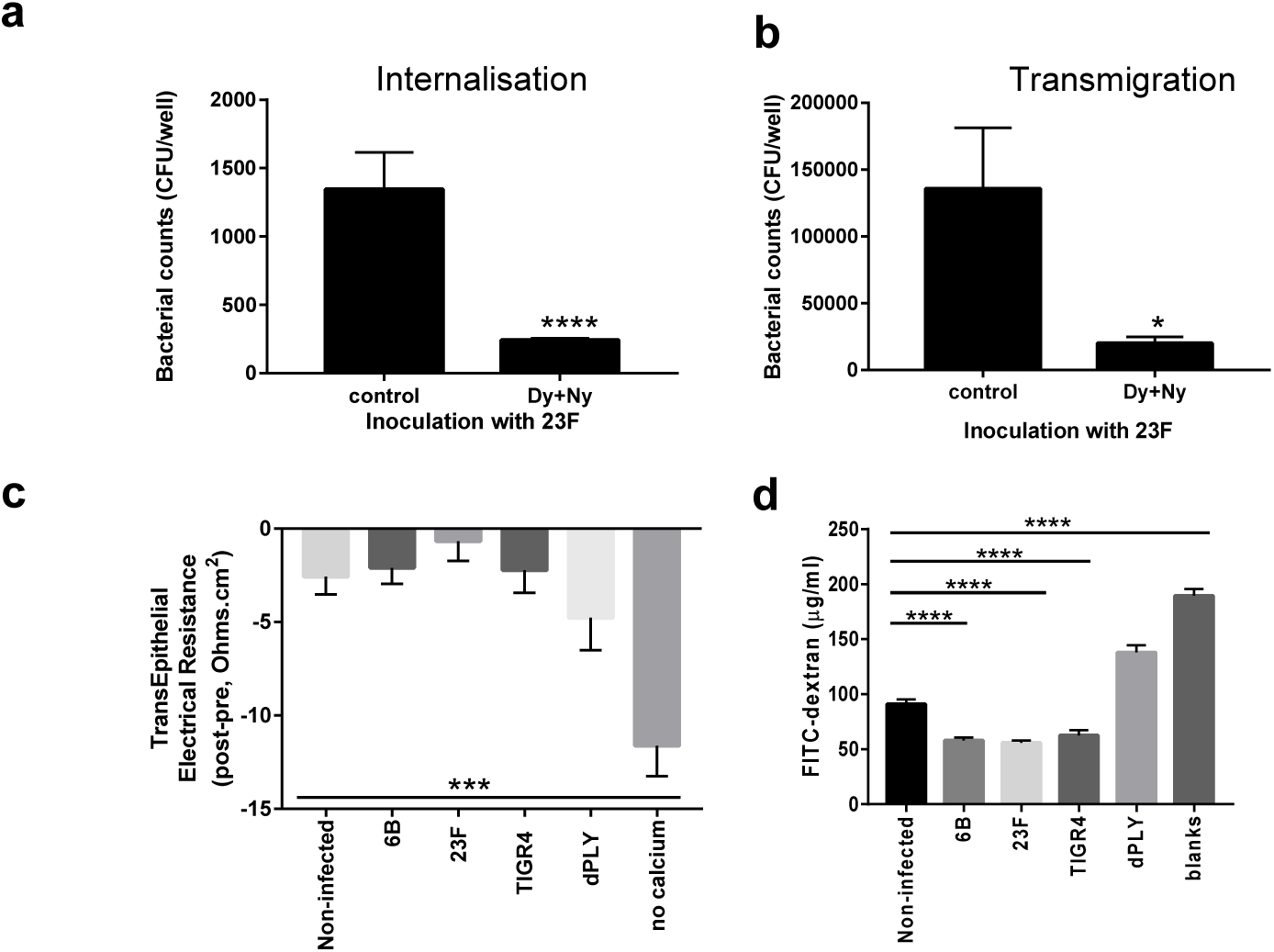
Transmigration of *S. pneumoniae* is endocytosis-dependent but does not disrupt barrier integrity or function. (a and b) Detroit 562 cells were treated with dynasore and nystatin for 30 minutes prior to and during incubation with 23F, for 1 hour. (a) Intracellular bacteria were quantified using a gentamicin protection assay and CFUs counted (****P = <0.0001, Mann-Whitney). (b) Transmigrated bacteria after one hour were counted by CFUs (*P = 0.013 Mann-Whitney). N = 4. (c and d) Cells were co-infected with pneumococci for three hours on transwell inserts and barrier function was measured. (c) Trans-Epithelial Electrical Resistance (TEER) was compared before and after exposure to pneumococci. N = 9. Calcium withdrawal was used as a positive control (***P = 0.0013, unpaired T-test, n = 3). (d) Permeability was assessed by leak to 4kDa FITC-dextran, collected from basal chamber. N = 4. ****P = <0.0001 comparing non-infected against pneumococci (Kruskall-Wallis). Blank inserts were used as a positive control (Mann-Whitney).

Together these data show the strain dependency of pneumococcal-epithelial interactions *in vitro* which mirror the epithelial micro-invasion (both internalisation and/ or transmigration of pneumococci) seen *in vivo*.

### Endocytosis-related micro-invasion and transmigration

To confirm that the pneumococcal endocytosis leads to transmigration across the epithelium, endocytosis was inhibited with dynasore and nystatin^16^. After one hour of infection, cellular uptake of 23F was inhibited by 82% (Figure 4a) and transmigration across the cell monolayer by 85% (Figure 4b). This inhibition of endocytosis and transmigration was also seen with TIGR4 (data not shown). In agreement with previous studies using A549 and Calu3 cells^30^, we found that viable pneumococci in Detroit 562 cells decreased over time (Supplementary Figure 5a). Using gentamicin treatment to eliminate the original extracellular inoculum, we observed *S. pneumoniae* in the apical and basal media (Supplementary Figure 5b), suggesting egress from the epithelium.

**Figure 5.**
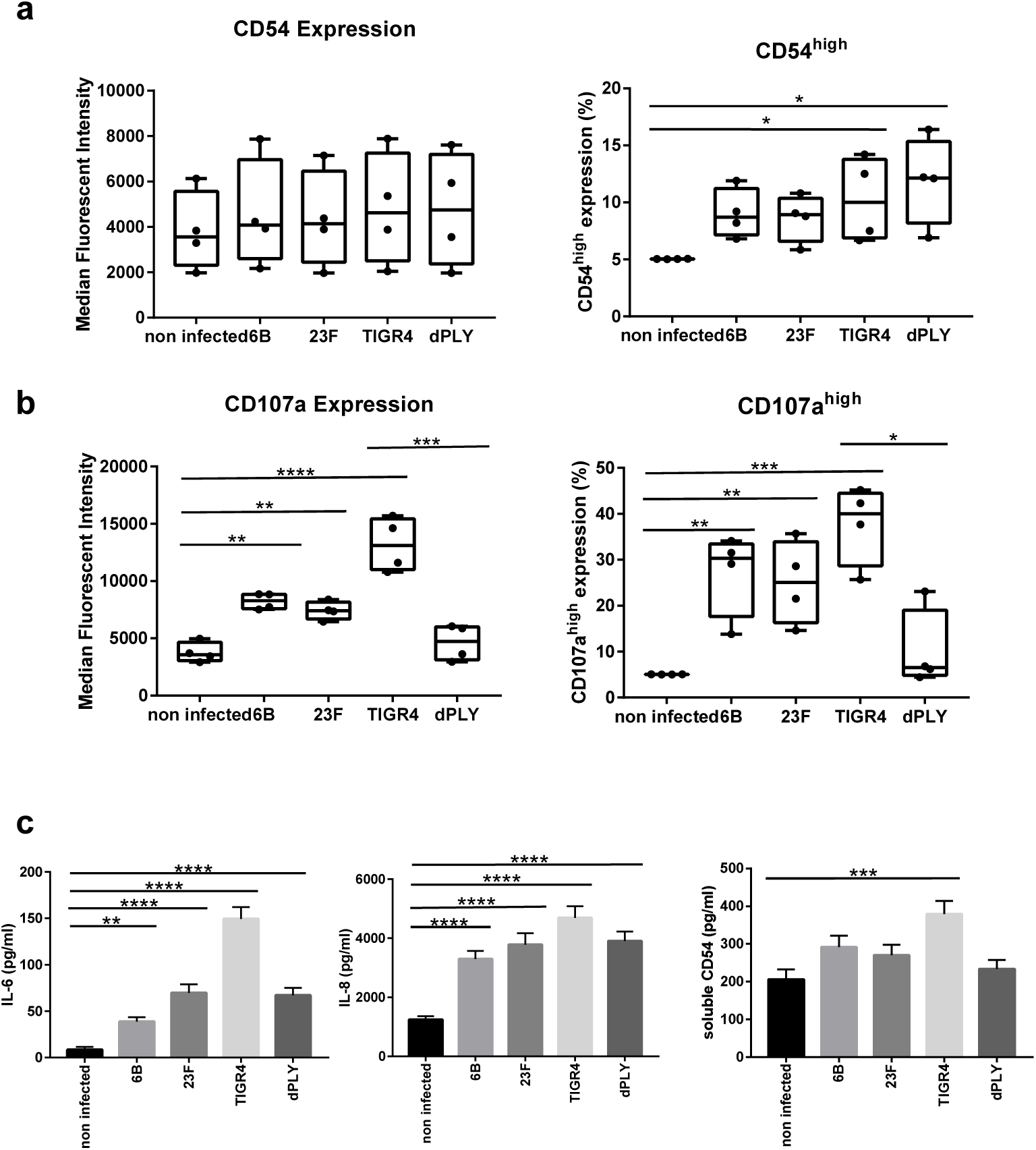
Epithelial CD107a and CD54 surface expression and secretion of IL-6, IL-8 and CD54 are upregulated after *S. pneumoniae* exposure *in vitro*. (a and b) Detroit 562 cells were stimulated with *S. pneumoniae* for 6 hours and the median fluorescence intensity and high-expressing cells for (a) CD54 and (b) CD107a were analysed by flow cytometry. n = 3 independent experiments (**** P < 0.0001 CD107a MFI, compared to non-infected cells, ANOVA. *** P = 0.0009 for MFI CD107a, TIGR4 v dPly, unpaired T-test. *** P < 0.001 CD107a high expression, compared to non-infected cells, ANOVA *P = 0.028 for high expression CD107a, TIGR4 vs. dPly, Mann-Whitney). (* P = 0.048 NI v TIGR4; * P = 0.0367 NI v dPly for CD54, ANOVA). (c) IL-6, IL-8 and CD54 in the supernatants from Detroit 562 cells stimulated with *S. pneumoniae* for 6 hours, were measured by ELISA. N = 6 independent experiments with replicates. (**** P <0.0001 IL-6; *** P = 0.0013 CD54; **** P < 0.0001 IL-8 (Kruskall-Wallis test)).

### Loss of epithelial cell barrier function is not a pre-requisite for micro-invasion by *S. pneumoniae*

*S. pneumoniae* has been shown to affect the integrity of epithelial barriers and tight junction function in murine models^13,31^. Having demonstrated pneumococcal micro-invasion without marked epithelial morphological disruption both by confocal and electron microscopy, we explored possible effects on epithelial barrier function by measuring Trans-Epithelial Electrical Resistance (TEER) and permeability to 4kDa FITC-dextran in Detroit 562 cells. TEER was not adversely affected by pneumococcal co-culture (Figure 4c) and permeability was maintained with all the pneumococcal strains at three hours post-infection (Figure 4d). Although co-localisation of pneumococci was seen with host proteins, we did not detect marked epithelial membrane carbohydrate or protein re-distribution (WGA, Claudin 4 JAM-A, β catenin, Supplementary Figure 3a). Similar observations were made with A549 cells and Calu3 cells (Supplementary Figure 4). These data suggest that pneumococcal micro-invasion was not dependent on loss of epithelial cell barrier function during early interactions.

### *S. pneumoniae* upregulates epithelial surface CD54 and CD107a *in vitro*

In a similar manner to primary cells collected from nasal curettes, we assessed Detroit 562 cell surface marker expression in response to *S. pneumoniae* (Supplementary Figure 6a-c). There were no significant changes in IL-22Ra1, HLADR or CD40 expression in response to 6B, 23F, TIGR4 and dPly-TIGR4 strains (Supplementary Figure 6d-f). However, CD54^high^ expression was significantly greater in response to the TIGR4 and dPly-TIGR4 strains, compared to non-infected cells (Figure 5a). Epithelial CD107a was upregulated in response to the 6B, 23F and TIGR4 strains (Figure 5b), but this was not seen with dPly-TIGR4. These data implicate pneumolysin in the induction of CD107a but not CD54 epithelial surface expression and support the possibility that CD107a is involved in pneumococcal-epithelial micro-invasion events seen *in vivo*.

**Figure 6.**
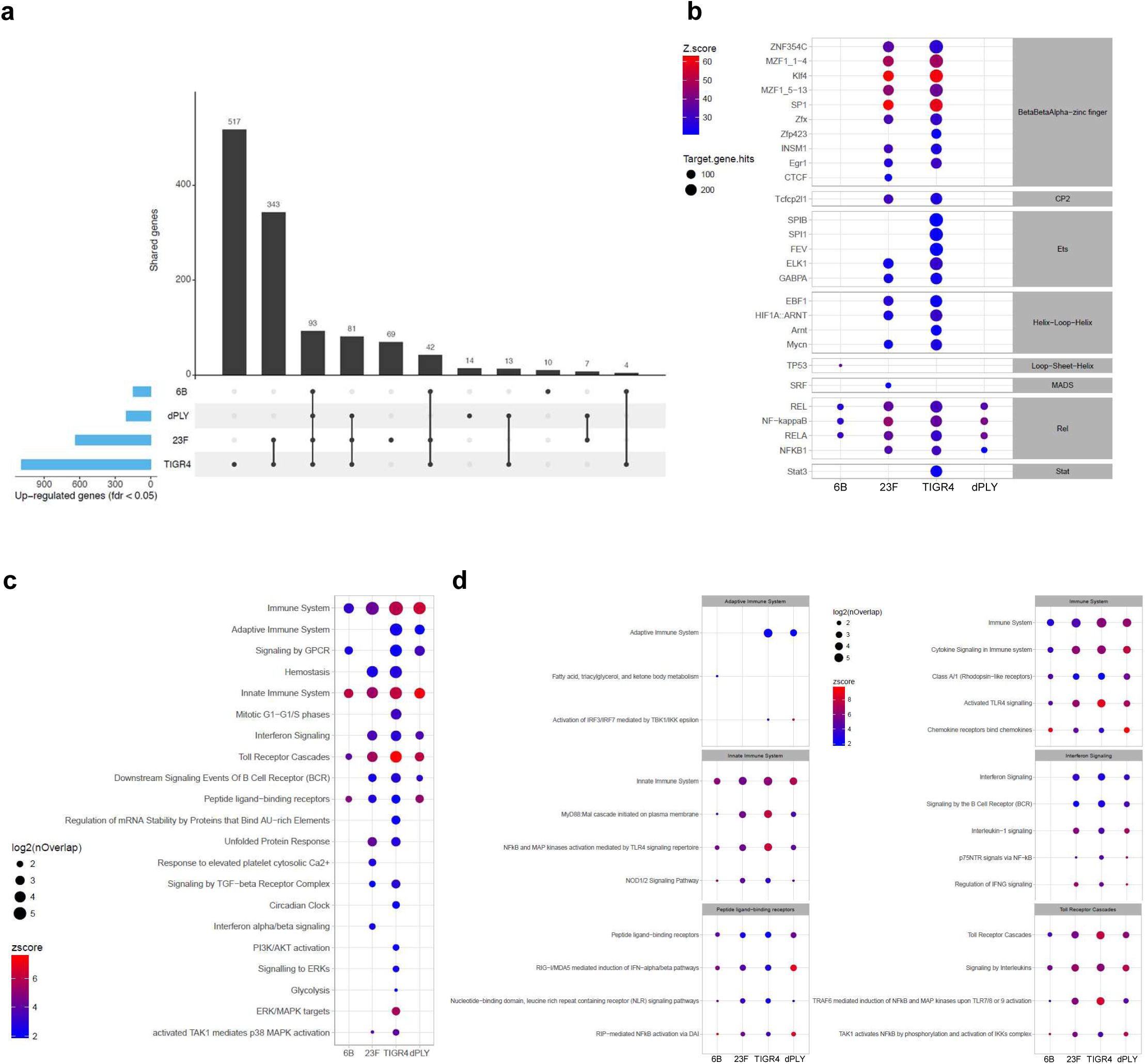
The epithelial transcriptomic response to *Streptococcus pneumoniae in vitro* is influenced by epithelial adhesion and micro-invasion. (a) The total number of Detroit 562 epithelial genes upregulated following exposure to 6B, 23F, TIGR4 and TIGR4-dPly pneumococci, compared to non-infected samples. N = 3 independent experiments with replicates. Genes with an FDR <0.05 were considered for further analysis. The matrix shows intersections for the four strains, sorted by size. Dark circles in the matrix indicates sets that are part of the intersection. (b) Bubble heat map showing the results of transcriptional factor analysis in Opossum for each strain. Transcriptional factors are ordered by family and then sorted by number of genes in the database. The colour represents the log2 z-scores (a continuum from blue (lowest values) to red (highest values) and the size shows the number of genes annotated to the particular pathway. (c) Bubble heat map showing the results of Reactome Pathway Analysis for upregulated genes in each strain compared to non-infected cells. Pathways were clustered in 20 groups using K-means on Jaccard index and the biggest pathway for each group was selected for representative annotation of each cluster. Individual pathways within clusters are shown in Supplementary Figure 8. The colour represents the log2 z-scores (a continuum from blue (lowest values) and red (highest values) and the size shows the number of genes annotated to each pathway. (d) Selected bubble heat map clusters representing the individual pathways within each cluster from (c).

### *Streptococcus pneumoniae* induces epithelial secretion of IL-6, IL-8 and CD54

To assess the inflammatory consequences of pneumococcal-epithelial infections *in vitro*, we measured a cytokine panel in Detroit 562 cell supernatants following *S. pneumoniae* incubation (Figure 5c). We detected a significant increase in IL-6 and IL-8, which were strain dependent and partially pneumolysin dependent. In line with surface marker observations, only TIGR4 significantly upregulated the secretion of soluble CD54, which was dependent on pneumolysin (Figure 5c). These data highlight the less inflammatory nature of the 6B strain compared to the more invasive strain, TIGR4, and the importance of pneumolysin in the epithelial response. Pneumococcal wild-type strain differences were not explained by differences in pneumolysin activity, as determined by hemolysis (Supplementary Figure 7).

**Figure 7.**
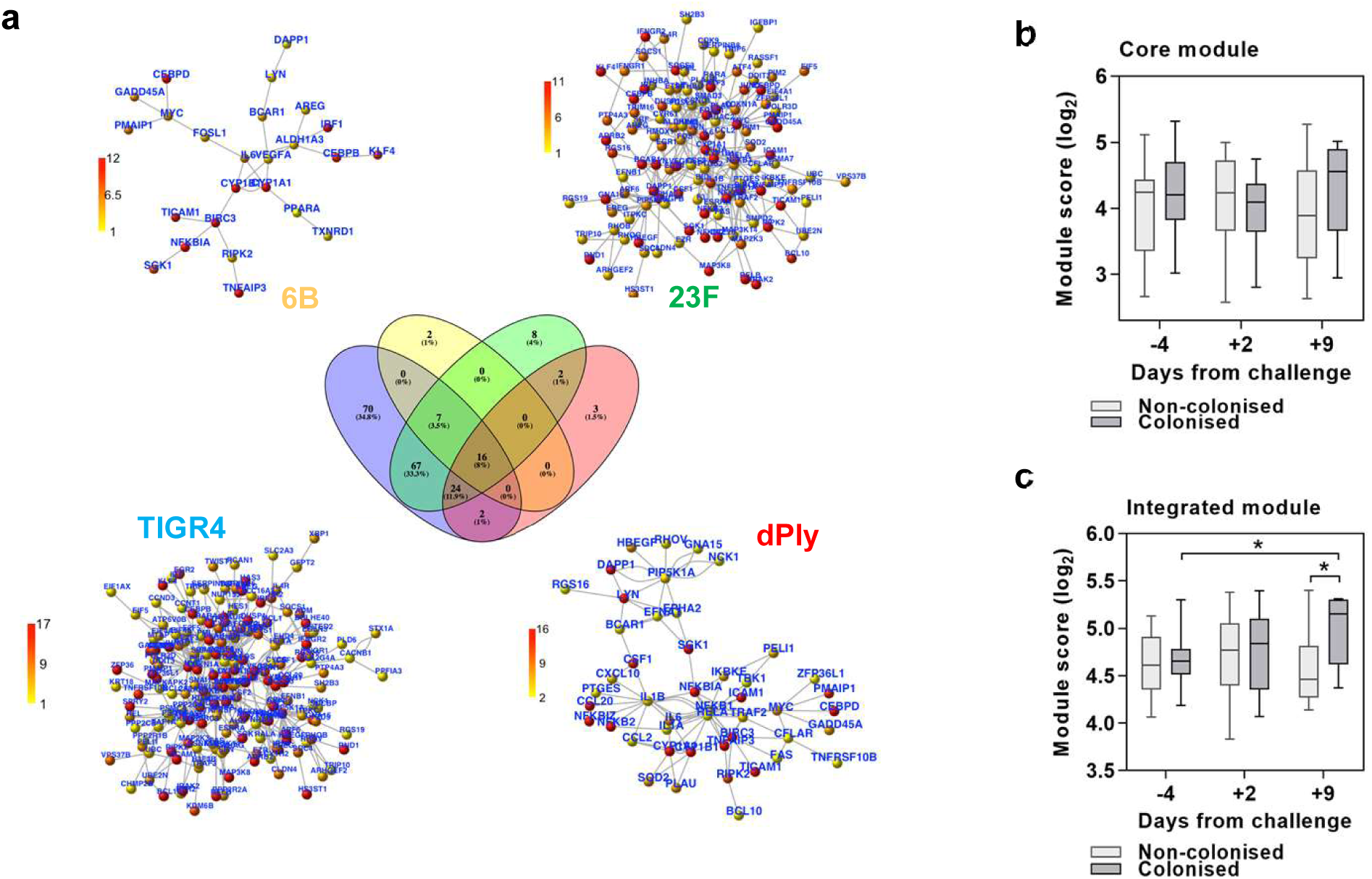
Epithelial-derived transcriptomic responses to *S. pneumoniae* in the EHPC model co-incides with bacterial clearance. (a) Network representation generated from the *in vitro* analyses for each strain. From the genes present in the networks, an integrated gene list (200 genes) and a core genes list (16 genes) were generated and tested as modules in the *in vivo* data. (b and c) Upregulated genes in comparison to the transcriptomes of carriage negative and carriage positive volunteers were compared between time points. Plots represent log2 TPM arithmetic mean for genes in the core (b) and integrated (c) modules. P = <0.05, (one tail Mann-Whitney test). Number of samples for carriage negative and positive, respectively: pre (17:14), day 2 (15:14), day 9 (7:7).

### Innate transcriptomic responses to epithelial infection by *S. pneumoniae in vitro*

To explore the hypothesis that the pattern of epithelial adhesion and micro-invasion results in differential epithelial responses, we performed RNAseq of Detroit 562 cells infected with our panel of *S. pneumoniae* strains. In comparison to non-infected epithelial cell cultures, infection with TIGR4 upregulated expression of 1127 genes (517 unique genes), 23F upregulated 650 genes (69 unique genes), and 6B upregulated 153 genes (10 unique genes), (Figure 6a). The pneumolysin deficient isogenic mutant of TIGR4 upregulated 220 genes (14 unique genes). Compared to non-infected cells, 1193 genes were upregulated by *S. pneumoniae* infection overall and 93 were core to all strains tested (differentially upregulated genes in Supplementary Excel file 1 and Reactome pathways in Supplementary Excel file 2).

To elucidate the pathways that drive the transcriptional responses to each strain, we also undertook transcription factor binding site enrichment analysis (Figure 6b, transcription factor binding site enrichment analysis in Supplementary Excel file 3). The responses to 6B and pneumolysin deficient strains were principally enriched in binding sites for the NFkB/Rel family of transcription factors, indicating that an NFkB activation pathway was the dominant driver for responses to these strains. In contrast, cellular responses to TIGR4 and 23F strains revealed enriched binding sites for more diverse transcription factors, suggesting broader molecular pathways by which these strains may influence cellular function. This analysis revealed particular enrichment of binding sites for beta-beta-alpha zinc finger superfamily of transcription factors, including KLF4, suggesting that these strains upregulate mitogen activated protein kinase pathways upstream of these transcription factors^32^.

The sets of genes upregulated by each strain individually were subjected to core and interactome and pathway-enrichment analysis for functional annotation of the cellular response at systems level (Figure 6c). From 20 clusters of pathways, significant enrichment of innate immune system responses was evident in all strains and included specific enrichment in Toll receptor cascades, peptide ligand-binding receptors, NFkB and MAP kinase activation, Interferon signalling and cytokine and chemokine signalling, (Figure 6c, 6d, Supplementary Figure 8). In general, each pneumococcal strain induced a distinct transcriptomic profile, with TIGR4 and 23F strains inducing a broader number of biological pathways than the 6B strain (Figure 6c). Fewer transcriptional responses at the level of functional pathways in response to dPly-TIGR4 is consistent with a role for pneumolysin in innate immune recognition of *S. pneumoniae*^33^. Fewer transcriptional responses to 6B is consistent with our observations that this strain exhibits the lowest level of cellular association and transmigration.

**Figure 8.**
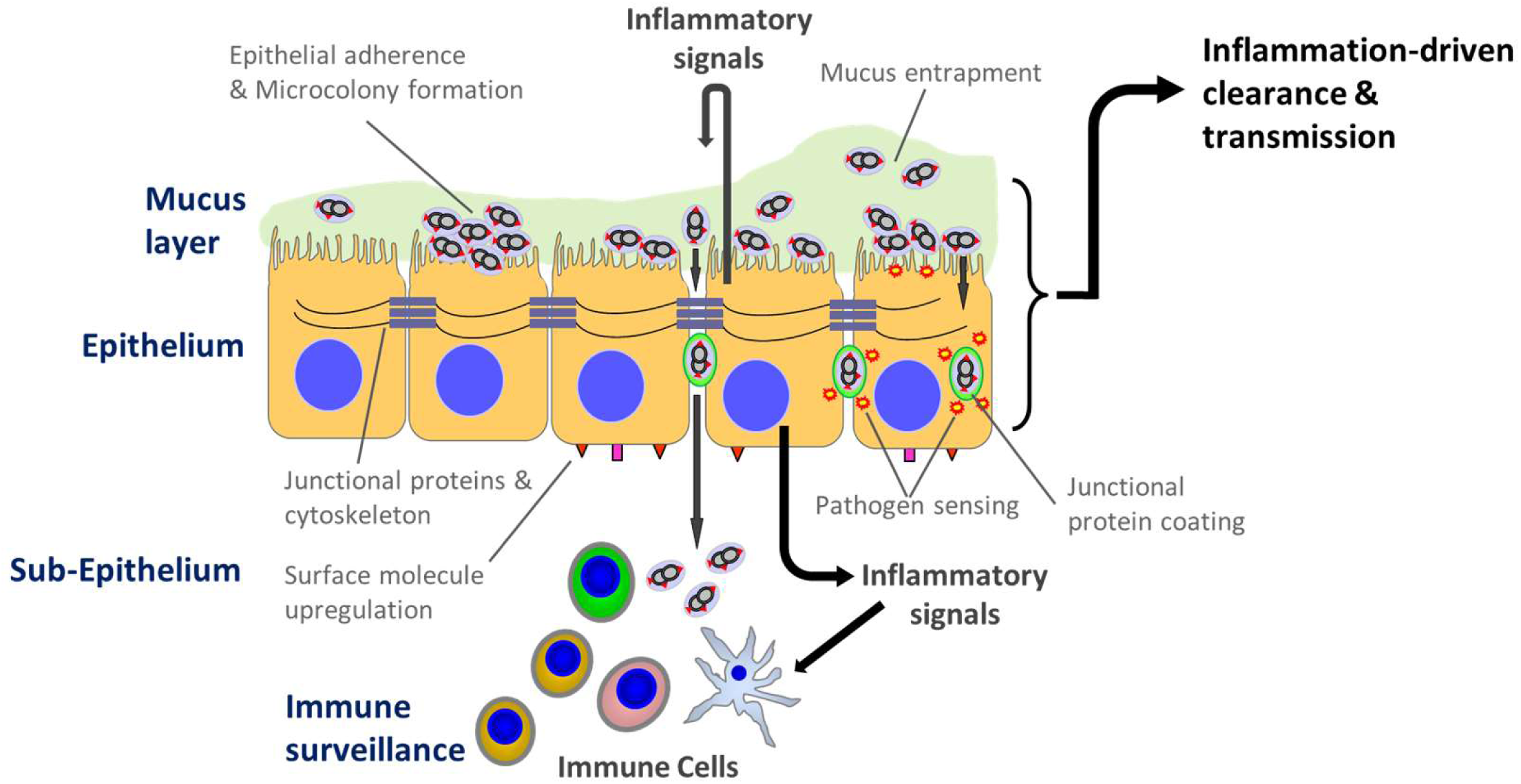
Model of pneumococcal colonisation at the human mucosal epithelium. Pneumococcal adhesion and micro-colony formation on the epithelial surface may lead to micro-invasion; internalisation of the bacteria and/ or transmigration across the epithelial barrier (micro-invasion). The epithelial-derived response is dependent on the subsequent pattern of interactions. Micro-invasion amplifies epithelial sensing and inflammation/ innate immunity, which we postulate leads to immune cell engagement. This process of epithelial sensing inflammation/ innate immunity may enhance both clearance and transmission. Co-association with junctional proteins may facilitate migration across the barrier.

### Epithelial transcriptomic responses to 6B are detectable *in vivo*

Next, we sought to test whether these epithelial transcriptomic responses to *S. pneumoniae* were also evident in the 6B EHPC model. We compared nasal transcriptomes sampled at two or nine days after inoculation to pre-inoculation samples. These revealed enrichment of 162 transcripts among individuals who became colonised (differential gene expression in Supplementary Excel file 4). These genes included Claudin 5 and Claudin 17, Defensin β 103A/B, Cadherin 16, Desmocollin 1 and Gap junction protein α1, which suggests cytoskeletal re-organisation two days post-inoculation in carriage positive individuals. By day 9 post-inoculation, molecules such as CCR3, matrix metalloproteinase 12 and MHC II molecules were enriched, suggesting activation of the nasal mucosa^34^. However, these genes were not significantly enriched for any immune response pathway annotation perhaps because of the statistical stringency necessary for genome-wide multiple testing and were not epithelial-specific.

Gene expression modules can successfully detect specific transcriptional programs in bulk tissue^35^. Therefore, as an alternative approach, we sought to target specific epithelial responses *in vivo* using the *in vitro* interactome. A direct comparison of transcripts for 6B revealed FOSL-1, an AP-1 transcription factor subunit, to be enriched with carriage, which has previously been reported to become activated following pneumococcal challenge in BEAS-2B and HEK293 cells^36^. To expand the sensitivity of the approach, we evaluated the expression of two transcriptional modules in the nasal samples (Figure 7). A ‘core’ module comprised of 16 interacting genes (Figure 7b) that were upregulated in the Detroit epithelium by all the pneumococcal strains from Figure 7a, and a second ‘integrated’ module that comprised of 200 interacting genes (Figure 7c), which were upregulated in the Detroit epithelium by at least one of the pneumococcal strains from Figure 7a. Among carriage negative individuals, we found no increase in expression in either gene expression modules (Figure 7b-c). In contrast among the individuals who became colonised, the increased expression of both modules, (significantly increased within the integrated module, Figure 7b,7c), suggests that the prominence of responses at day nine may reflect time-dependent proliferation of the 6B bacterial inoculum, epithelial engagement and the subsequent epithelial activation. These responses coincide around the time of bacterial clearance.

## DISCUSSION

It is widely accepted that colonisation of the URT mucosa by many potentially pathogenic bacterial species, involves transient association with the overlying mucus layer but that adherence to the epithelial surface avoids mucus entrapment and muco-ciliary clearance^14,37^. While evading immunity, bacterial replication then occurs prior to onward transmission to a new host. Even for *S. pneumoniae*, the mechanisms underlying this transition from acquisition to established colonisation, and then transmission and/or disease, are not well understood. By combining an EHPC model and *in vitro* human cell culture systems, we show that pneumococcal colonisation leads to epithelial adherence, micro-colony formation and migration across the epithelial barrier without disease, which we have termed micro-invasion. The finding of co-association with junctional proteins provides a possible mechanism for micro-invasion. Clearance of the pneumococcus from the mucosa *in vivo* occurs in the context of micro-invasion. Indeed, epithelial micro-invasion and not bacterial load *per s*e, appears to dictate unique transcription factor binding site signatures, downstream cytokine, chemokine and metabolic pathway enrichment (Figure 8).

The concept of mucosal micro-invasion moves us away from current models of colonisation where the pneumococcus is held up at the epithelial surface and that invasion leads to disease. Our data are supported by murine colonisation experiments^38,39^ and the detection of pneumococcal DNA in blood of otherwise healthy, colonised children^40^. In line with other cell culture and murine models^13,16,38,41^, we have demonstrated that pneumococcal micro-invasion occurs by endocytosis through the formation of cytoplasmic vacuoles, and by paracellular transcytosis across the epithelium. This process appears to be association with the formation of epithelial folds different from the membrane ruffles previously reported with conventionally intracellular pathogens such as *Salmonella* typhimurium^42^. Our study emphasises the active nature of these processes without marked cellular damage or loss of barrier function. Micro-invasion of the epithelium may overcome the inaccessibility of PAMP receptors located either at the epithelial basolateral surface or intracellularly^43,44^. We have found that at early time points, micro-invasion *in vitro* is also characterised by pneumococcal co-association with JAM-A and β catenin without evidence of barrier dysfunction. In murine models at later time points, TLR-dependent Claudin 7 and Claudin 10 down-regulation enhanced pneumococcal translocation across the epithelium has been observed^13^. While this may occur as inflammation becomes more prominent, our data suggest that micro-invasion can happen without barrier disruption and that the epithelium plays an active role in the regulation of these complex host-pathogen interactions^12^. How micro-invasion relates to the risk of invasive pneumococcal disease remains to be determined but as demonstrated by the EHPC model, micro-invasion may occur in healthy individuals without symptoms.

The differential impact of micro-invasion on the epithelial transcriptomic response was striking with considerable enrichment of multiple diverse transcription factors and signalling pathways related to innate immunity with the most invasive pneumococcal strains. The interactome module highlights that although enrichment of gene activation in response to all pneumococcal strains is apparent (the core module), a more diverse range of epithelial gene activation occurs between strains, reflecting specific host-pathogen interactions. Nasal colonisation in murine models is proinflammatory^45,46^, but our *in vitro* and EHPC experiments with serotype 6B suggests this is not always the case, whereby pneumococcal-host cell adherence, growth, endocytosis and paracellular migration can be established without a marked host response. In the EHPC, clearance occurred around the time the epithelial transcriptomic response was most prominent. Weiser and colleagues argue that the colonising pneumococcus induces a host inflammatory response that mediates clearance, but also promotes nutrient acquisition, mucus production and onward transmission^14,18,47^, which is supported by murine models^48^ and epidemiological studies of viral-coinfection^49^. It has been suggested that pneumolysin induces neutrophil influx and degranulation, leading to increased secretions from the nasopharynx, so promoting transmission^5,18^. We therefore suggest that the epithelial innate response resulting from pneumococcal colonisation could promote both clearance and transmission.

The enrichment of binding sites for the transcription factor KLF4 transcription factors seen in our *in vitro* data suggests a counter-regulatory process aimed at minimising inflammation. KLF4 plays a role in barrier function, suppression of NFκB-dependent IL-8 secretion, regulation of IL-10 expression via TLR9-MyD88 and Yes-1, in response to *S. pneumoniae*, which is partially dependent on autolysin LytA^32,50,51^. Thus, by exploiting micro-invasion, the pneumococcus may carefully calibrate the host innate immune/ inflammatory response to promote survival through transmission.

A range of pneumococcal PAMPs including pneumolysin, may trigger this epithelial sensing process^18,52^. We found pneumolysin to be a prominent inducer of epithelial surface molecule upregulation, cytokine production and transcriptomic inflammatory response *in vitro*. Following internalisation of the pneumococcus by neutrophils, pneumolysin has been shown to induce ROS following bacterial autolysis, which leads to cellular activation^53^. We speculate that pneumolysin released intracellularly may signal directly in epithelial cells or through host cell pore-formation leading to entry of other PAMPs. Mediated by autolysin, the pneumococcus undergoes autolysis during stationery growth phase, resulting in additional PAMPs’ release including bacterial DNA^53^. In mice, pneumococcal DNA triggers inflammation through a DAI/STING/TBK1/IRF3 cascade^45,54^, a response that may contribute to pneumococcal clearance. In our *in vitro* transcriptomic analyses, we detected enriched activation of IRF3/IFR7 in dPly-TIGR4. We suggest that in the context of micro-invasion, pneumococcal DNA may also act as a trigger of epithelial sensing, inducing inflammation^54^.

Our findings are limited by the pneumococcal strains available for testing in the EHPC model to enable direct comparisons with *in vitro* data. Nonetheless, the use of different strains *in vitro* has enabled us and others^55,56^ to determine the impact of different patterns of epithelial adherence and invasion on the host inflammatory/ innate immune response. Currently, the EHPC model does not allow us to definitively assess human to human transmission although the appropriate methods are under development.

In conclusion, our data implicates the novel finding of micro-invasion during pneumococcal colonisation of otherwise healthy humans, promoting epithelial-derived innate immunity/ inflammation and ultimately clearance. The pathways critical for onward pneumococcal transmission remain to be determined in humans but based on murine models, we propose that this epithelial response may also facilitate transmission. The balance between innate immunity/ inflammation-driven transmission and clearance may be further modulated by the frequency of carriage events, pneumococcal strain co-colonisation, viral co-infections and other environmental pressures^1,22,48^. Our approach combining *in vitro* with *in vivo* human systems offers a potentially tractable way to further interrogate these processes.

## ACKNOWLEDGEMENTS

This study was funded by the Wellcome Trust (Grant 106846/Z/15/Z). DF is supported by the Medical Research Council (grant MR/M011569/1), Bill and Melinda Gates Foundation (grant OPP1117728) and the National Institute for Health Research (NIHR) Local Comprehensive Research Network. The authors wish to thank the EHPC Clinical Team at LSTM and all the volunteers. Confocal imaging facilities at LSTM were funded by a Wellcome Trust Multi-User Equipment Grant (104936/Z/14/Z). Flow cytometric acquisition at LSTM was funded by a Wellcome Trust Multi-User Equipment Grant (104936/Z/14/Z). LytA PCR was performed by Prof. D Bogaert, University of Edinburgh, UK. For *in vitro* data, RNAseq library preparation was undertaken at UCL through the UCL/UCLH Biomedical Research Centre and MRC funded Pathogen Genomics Unit (G0900950). For the *in vivo* data, RNAseq library preparation was performed and by Carl Anderson at the Wellcome Sanger Institute, Hinxton. EM was performed by Dr. Mark Turmaine in the Biosciences EM facility at UCL.

## AUTHOR CONTRIBUTIONS

CMW, SPJ, DMF and RSH conceived and designed the study. CMW, SP, SPJ, JR, EN, CS acquired the data. CMW, CV, SPJ, MN, JSB, DMF, RSH analysed and interpreted the data. CMW wrote the first draft of the manuscript. CMW, CV, SP, SPJ, JR, EN, CS, MN, JSB, DMF, RSH commented on and approved the manuscript.

## METHODS

### Bacteria

*S. pneumoniae* clinical strains used were 6B (BHN 418^57^), 23F (P1121^58^) and TIGR4 (P1672^59^), together with a pneumolysin deficient TIGR4 mutant strain (P1672, a kind gift from Prof T Mitchell, University of Birmingham). Stocks of bacterial aliquots grown to O.D._600nm_ 0.3 were stored at −80°C, defrosted, resuspended in cell culture media and used once. Colony forming units (CFU) were counted on horse blood agar plates (EO Labs).

### Experimental Human Pneumococcal Carriage Model (EHPC)

Following written informed consent, healthy non-smoking adults between the ages of 18 – 59 were inoculated with 80,000 CFU/nostril live 6B *S. pneumoniae* (BHN418), grown to mid-log phase in vegetone broth as previously described^60^. Individuals naturally colonised with *S. pneumoniae* or in regular contact with at-risk individuals were excluded (except for EM images in Supplementary Figure 1b). Blood and mucosal T cell and antibody immunity were not systematically measured in this cohort but previous studies of volunteers from this population have demonstrated background T cell and antibody-mediated anti-pneumococcal immunity, but a relationship between pre-challenge immunity and the development of carriage in the model has not been established^21,23^. To exclude the influence of viral-coinfection, EHPC model volunteers who had coryzal or flu-like symptoms prior to challenge were excluded. Volunteers were screened for the presence of 20 common respiratory viruses before inoculation. 1/13 asymptomatic volunteers (volunteer 1) was positive for enterovirus. Nasal washes and mucosal cells (curette biopsy) from the inferior turbinate were obtained by PBS syringe and curettage using a plastic Rhino-probe™ (Arlington Scientific), in this order of collection respectively, before pneumococcal inoculation. Sampling was then repeated on days 2, 6, 9, and 14 – 27 post pneumococcal inoculation^61^. Volunteers received the TIV vaccine intramuscularly at day 3 post pneumococcal inoculation. Bacteria collected from nasal washes were quantified by CFU counts. At each time-point, two curettage samples were obtained and processed for confocal immunofluorescence, flow cytometry, primary cell culture and /or transcriptomic analysis by RNA sequencing (RNAseq). There were no adverse events. The carriage status of each volunteer was blinded until sample analysis was completed.

Ethical approval was given by NHS Research and Ethics Committee (REC)/Liverpool School of Tropical Medicine (LSTM) REC, reference numbers: 15/NW/0146 and 14/NW/1460 and Human Tissue Authority licensing number 12548.

### Human Respiratory Tract Epithelial Cell Lines

Human pharyngeal carcinoma Detroit 562 epithelial cells (ATCC_CCL-138) and human bronchial carcinoma Calu3 epithelial cells (ATCC_HTB-55) were grown in 10% FCS in alpha MEM media (Gibco). Human alveolar epithelial carcinoma A549 epithelial cells (ATCC_CCL-185) were grown in 10% FCS with 1% L-glutamine in Hams/F-12 media (Gibco).

### Pneumococcal-epithelial cell co-culture

#### Association and Invasion assays

confluent Detroit 562 (typically day 8 post plating), Calu3 (typically day 10 post plating) and A549 (typically day 4 post plating) monolayers cultured on 12 well plates (Corning), were exposed to *S. pneumoniae* for three hours in 1% FCS alpha MEM (∼MOI approximately 1 cell: 10 pneumococci). The medium was removed, and cells washed three times in HBSS^+/+^. Cells were incubated in 1% saponin for 10 minutes at 37°C and lysed by repetitive pipetting. Dilutions of bacteria were plated on blood agar and colonies counted after 16 hours. To quantify internalised bacteria, 100μg/ml gentamicin was added for 1 hour to the cells, which were washed another three times, before incubating with 1% Saponin and plating on blood agar plates. CFUs were counted after 16 hours incubation at 37°C, 5% CO_2_. There were no differences in pneumococcal pre-or post-inoculum, or growth density between the strains, in the cell supernatant three hours post-infection.

#### Transmigration assay

Detroit 562 cells were cultured on 3μm pore, PET Transwell Inserts (ThermoFisher) typically for 10 days to achieve confluent, polarised monolayers. Calu3 cells were plated onto Transwell inserts typically for 12 days and A549 cells typically for 6 days. Cell culture media was changed 1 hour prior to addition of bacteria to 1% FCS (250μl apical chamber, 1ml basal chamber). Resistance was recorded before and after *S. pneumoniae* were added using an EVOM2 (World Precision Instruments). 1mg/ml FITC-dextran (Sigma Aldrich) was added to the apical chamber of selected inserts to assess permeability. Approximately 12 million bacteria were added to the cells (∼MOI 1 cell: 25 bacteria). During the time course, 50μl was removed, diluted and plated, from the basal chamber to measure bacterial load by counting CFU/well. Permeability was recorded using a FLUOstar Omega (BMG Labtech) at 488nm.

#### Endocytosis inhibition assays

Detroit 562 cells cultured on 12 well plates were treated with 100μM dynasore (Cambridge Biosciences) and 7.5μg/ml nystatin (Sigma Aldrich) to block endocytosis for 30 minutes prior to, and for the duration of pneumococcal infection incubation period. DMSO was used as a control. Cells were washed and treated with gentamicin and lysed with saponin as described above.

### Confocal Microscopy

For the *in vivo* analysis, mucosal cells derived by curettage from the EHPC model were placed directly into 4% PFA for 1 hour. Cells were cytospun onto microscope slides and allowed to air dry. For the *in vitro* analysis, epithelial cell lines on transwell membranes were fixed in either 4% PFA (Pierce, Methanol Free) or 1:1 mix of methanol: acetone for 20 minutes. For both *in vivo* and *in vitro* samples, cells were permeabilised with 0.2% Triton X-100 for 10 minutes and blocked for 1 hour in blocking buffer (3% goat serum and 3% BSA in PBS) before incubation with either pneumococcal antisera Pool Q (for 6B or 23F detection, SSI Diagnostica), or pneumococcal antisera Type 4 (for TIGR4 and dPly detection, SSI Diagnostica) to detect pneumococci, and JAM-A, Claudin-4 or β-catenin primary antibodies to detect cellular junctional proteins (see Supplementary Information) for one hour and then secondary and/or conjugated antibodies for 45 minutes. DAPI solution was added for 5 minutes. After washing, the stained samples were mounted using Aqua PolyMount (VWR International) with a coverslip onto a microslide. The entire cytospin for each sample was manually viewed by microscopy for detection of pneumococci. Multiple fields of view (>3) were imaged for each transwell insert, for each condition. Images were captured using either an inverted LSM 700, LSM 880, or TissueFAXS Zeiss Confocal Microscope. Z stacks were recorded at 1μm intervals at either 40x oil or 63x oil objectives. For the *in vivo* samples, the operator (CMW) was blinded to the colonisation status of the volunteer at the time of sampling.

### Electron Microscopy

Detroit 562 cells cultured on either transwells (scanning EM) or coverslips (transmission EM) were fixed with 2% paraformaldehyde and 1.5% glutaraldehyde in 0.1 M cacodylate buffer and post-fixed in 1% OsO4 / 1.5% K4Fe(CN)6 in 0.1 M cacodylate buffer pH7.3. Samples were dehydrated in graded ethanol-water series and infiltrated with Agar 100 resin mix. Ultra-thin sections were cut at 70-80 nm using a diamond knife on a Reichert ultracut microtome. Sections were collected on 300 mesh copper grids and stained with lead citrate. Samples were viewed in a Joel 1010 transition electron microscope and images recorded using a Gatan Orius camera. Further details of TEM and SEM methods can be found in the Supplementary information.

### Flow cytometry

For the *in vivo* analysis, cells from two curette samples were incubated in cold PBS++ (PBS supplemented with 5mM EDTA and 0.5% FCS) were dislodged by pipetting and centrifuged at 440g for 5 mins at 4°C. Supernatant was removed, and cells resuspended in 25μl of PBS++ with Live/Dead™ Fixable Violet Dead Cell Stain (ThermoFisher). After 15 minutes incubation on ice, an antibody cocktail to stain for epithelial surface marker expression (see Supplementary Information) was added and incubated for another 15 minutes. Samples were vortexed, resuspended in 3.5mls of PBS++ and filtered over a pre-wetted 70μm filter. Samples were transferred to a 5ml FACS tube, centrifuged and resuspended in 200μl Cell Fix (BD Biosciences) and acquired on LSRII Flow Cytometer (BD Biosciences). Compensation was run and applied for each experimental replicate and voltages consistent throughout. Isotype controls (Biolegend) and FL-1 controls were also run for each antibody. Analyses of data was performed using FlowJo LLC version 10 software. Data was performed on the gated epithelial cell population and only samples containing 500 or more cells are reported.

For the *in vitro* analysis, confluent monolayers of Detroit 562 cells on 6 well plates were incubated with *S. pneumoniae* for 6 hours in 1% FCS phenol free alpha MEM (base media, Life Technologies). Cells were washed three times in PBS and gently lifted from the plate using a cell scraper in 300μl of base media supplemented with 1mM EDTA. Samples were transferred to 5ml FACS tubes and placed on ice for the duration of the protocol. Each cell sample was incubated with an antibody cocktail for epithelial surface marker expression (see Supplemental Information) and Live/Dead™ Fixable Zombie Blue UV Cell Stain (Biolegend) for 30 minutes before rinsing in 1ml base media and centrifuging at 300g for 5 minutes at 4°C. Cells were fixed in 600μl of 4% PFA and acquired on an LSR II Flow Cytometer (BD Biosciences). Compensation was run and applied for each experimental replicate and voltages consistent throughout. Isotype controls (BD Biosciences), FL-1 and single stains were also run for each experiment. Samples were acquired until 300,000 events had been collected. Analyses were performed using FlowJo LLC version 10 software.

### ELISAs

Supernatant from Detroit 562 cells that had been incubated with *S. pneumoniae* for 6 hours, was collected for cytokine analysis. IL-6, IL-8 and ICAM-1 DuoSet® ELISA kits were purchased from R&D Systems and protocol followed according to manufacturers’ instructions.

### RNA samples and sequencing (RNASeq)

Mucosal curettage samples and epithelial cell cultures (incubated with or without *S. pneumoniae* for 3 hours) were collected in RNALater (ThermoFisher) and stored at −80°C. RNA Extraction was performed using the RNEasy micro kit (Qiagen) and genomic DNA was removed with on column DNA digestion or with the Turbo DNA-free Kit (Qiagen). Extracted RNA quality was assessed and quantified using a BioAnalyser (Agilent 2100). For mucosal curettage samples, library preparation and RNA-sequencing (Illumina Hiseq 4000, 15 lanes in total with 7 samples per lane, 100bp and 75bp paired-end reads) were performed at the Beijing Genome Institute (China) or the Wellcome Sanger Institute (UK), respectively. For the Detroit 562 samples, library preparation was performed using the Kappa Hyperprep kit (Roche Diagnostics) and sequencing was undertaken by the Pathogens Genomic Unit (UCL) on the Illumina Nextseq using the Nextseq 500/550 High Output 75 cycle kit (Illumina) according to manufacturers’ instructions, giving 15–20 million 41bp paired end reads per sample.

All data processing and analysis was conducted in R, language and environment for statistical computing (https://www.R-project.org). Paired end reads were mapped to the Ensembl human transcriptome reference sequence (homo sapiens GRCh38). Mapping and generation of read counts per transcript were performed using Kallisto^62^, based on pseudoalignment. R/Bioconductor package Tximport was used to import the mapped counts data and summarise the transcripts-level data into gene level data^63^. DESeq2 and SARTools packages^64^, were used for differential gene expression analysis, using a false discovery rate (FDR) <0.05, following normalisation with a negative binomial generalised linear model. In addition, transcript abundance for protein coding genes was expressed as log_2_-transformed transcripts per million (TPM) by normalising the read counts to gene length and then read depth. Transcriptomes from nasal samples ranged from 16 to 66 million reads per sample and in principle component analysis, revealed a batch effect that was corrected by the Combat function in the Surrogate Variable Analysis (SVA) R/Bioconductor package^65^.

Identification of interacting networks and pathway enrichment analysis was performed using XGR R package^66^. Interacting genes (supported by at least one experimental source of evidence) were identified from the Pathway Commons database for directed interactions. These were then subjected to Reactome pathway enrichment analysis with FDR<0.05. K-means clustering of Jaccard indices to quantify similarity between the composition of genes mapping to each pathway was used to identify 20 groups of pathways, from which the pathway consisting of the largest total number of genes was selected to provide a representative annotation for each group. Transcription factor binding site enrichment analysis was performed using the human single site analysis function in oPOSSUM^67^. The expression of gene modules in the nasal transcriptomes was represented by the mean Log_2_ TPM value of the genes in each module.

### Code and RNAseq data availability

Codes used to process the RNAseq data can be found at https://github.com/cristina86cristina. RNAseq data from the EHPC are accessible through GEO Series Number GSE124949. RNAseq data from the Detroit 562 cell infections is accessible through the ArrayExpress database at EMBL-EBI under accession number E-MTAB-7841. Data from differential gene expression from upregulated genes following infection, Reactome pathways and transcription factor analysis can be found in Supplemental Excel Files 1 – 4.

### Statistics

All experiments were conducted with replicates in three or more independent experiments unless stated otherwise. Error bars represent SEM unless stated otherwise. GraphPad Prism Version 10 was used to perform two-tailed parametric (t-tests or ANOVA) or non-parametric (Mann-Whitney or Kruskal-Wallis tests) analysis, which was based on the Shapiro-Wilk normality test. P values less than 0.05 were considered significant and are derived from non-parametric statistical tests unless stated otherwise.

## REFERENCES

1. O’Brien, K. L. et al. Burden of disease caused by Streptococcus pneumoniae in children younger than 5 years: global estimates. Lancet 374, 893–902, doi:10.1016/S0140-6736(09)61204-6 (2009).

2. Whitney, C. G. et al. Decline in invasive pneumococcal disease after the introduction of protein-polysaccharide conjugate vaccine. N Engl J Med 348, 1737–1746, doi:10.1056/NEJMoa022823 (2003).

3. Hanage, W. P. et al. Carried pneumococci in Massachusetts children: the contribution of clonal expansion and serotype switching. Pediatr Infect Dis J 30, 302–308, doi:10.1097/INF.0b013e318201a154 (2011).

4. Jochems, S. P., Weiser, J. N., Malley, R. & Ferreira, D. M. The immunological mechanisms that control pneumococcal carriage. PLoS Pathog 13, e1006665, doi:10.1371/journal.ppat.1006665 (2017).

5. Diavatopoulos, D. A. et al. Influenza A virus facilitates Streptococcus pneumoniae transmission and disease. FASEB J 24, 1789–1798, doi:10.1096/fj.09-146779 (2010).

6. Pido-Lopez, J., Kwok, W. W., Mitchell, T. J., Heyderman, R. S. & Williams, N. A. Acquisition of pneumococci specific effector and regulatory Cd4+ T cells localising within human upper respiratory-tract mucosal lymphoid tissue. PLoS Pathog 7, e1002396, doi:10.1371/journal.ppat.1002396 (2011).

7. Zhang, Q. et al. Characterisation of regulatory T cells in nasal associated lymphoid tissue in children: relationships with pneumococcal colonization. PLoS Pathog 7, e1002175, doi:10.1371/journal.ppat.1002175 (2011).

8. Pennington, S. H. et al. Polysaccharide-Specific Memory B Cells Predict Protection against Experimental Human Pneumococcal Carriage. Am J Respir Crit Care Med 194, 1523–1531, doi:10.1164/rccm.201512-2467OC (2016).

9. Mitsi, E. et al. Agglutination by anti-capsular polysaccharide antibody is associated with protection against experimental human pneumococcal carriage. Mucosal Immunol 10, 385–394, doi:10.1038/mi.2016.71 (2017).

10. Ratner, A. J., Lysenko, E. S., Paul, M. N. & Weiser, J. N. Synergistic proinflammatory responses induced by polymicrobial colonization of epithelial surfaces. Proc Natl Acad Sci U S A 102, 3429–3434, doi:10.1073/pnas.0500599102 (2005).

11. Eisele, N. A. & Anderson, D. M. Host Defense and the Airway Epithelium: Frontline Responses That Protect against Bacterial Invasion and Pneumonia. J Pathog 2011, 249802, doi:10.4061/2011/249802 (2011).

12. Whitsett, J. A. & Alenghat, T. Respiratory epithelial cells orchestrate pulmonary innate immunity. Nat Immunol 16, 27–35, doi:10.1038/ni.3045 (2015).

13. Clarke, T. B., Francella, N., Huegel, A. & Weiser, J. N. Invasive bacterial pathogens exploit TLR-mediated downregulation of tight junction components to facilitate translocation across the epithelium. Cell Host Microbe 9, 404–414, doi:10.1016/j.chom.2011.04.012 (2011).

14. Weiser, J. N., Ferreira, D. M. & Paton, J. C. Streptococcus pneumoniae: transmission, colonization and invasion. Nat Rev Microbiol 16, 355–367, doi:10.1038/s41579-018-0001-8 (2018).

15. Agarwal, V., Asmat, T. M., Dierdorf, N. I., Hauck, C. R. & Hammerschmidt, S. Polymeric immunoglobulin receptor-mediated invasion of Streptococcus pneumoniae into host cells requires a coordinate signaling of SRC family of protein-tyrosine kinases, ERK, and c-Jun N-terminal kinase. J Biol Chem 285, 35615–35623, doi:10.1074/jbc.M110.172999 (2010).

16. Asmat, T. M., Agarwal, V., Saleh, M. & Hammerschmidt, S. Endocytosis of Streptococcus pneumoniae via the polymeric immunoglobulin receptor of epithelial cells relies on clathrin and caveolin dependent mechanisms. Int J Med Microbiol 304, 1233–1246, doi:10.1016/j.ijmm.2014.10.001 (2014).

17. Ratner, A. J., Aguilar, J. L., Shchepetov, M., Lysenko, E. S. & Weiser, J. N. Nod1 mediates cytoplasmic sensing of combinations of extracellular bacteria. Cell Microbiol 9, 1343–1351, doi:10.1111/j.1462-5822.2006.00878.x (2007).

18. Zafar, M. A., Wang, Y., Hamaguchi, S. & Weiser, J. N. Host-to-Host Transmission of Streptococcus pneumoniae Is Driven by Its Inflammatory Toxin, Pneumolysin. Cell Host Microbe 21, 73–83, doi:10.1016/j.chom.2016.12.005 (2017).

19. Malley, R. et al. Recognition of pneumolysin by Toll-like receptor 4 confers resistance to pneumococcal infection. Proc Natl Acad Sci U S A 100, 1966–1971 (2003).

20. Ratner, A. J. et al. Epithelial cells are sensitive detectors of bacterial pore-forming toxins. J Biol Chem 281, 12994–12998, doi:10.1074/jbc.M511431200 (2006).

21. Ferreira, D. M. et al. Controlled human infection and rechallenge with Streptococcus pneumoniae reveals the protective efficacy of carriage in healthy adults. Am J Respir Crit Care Med 187, 855–864, doi:10.1164/rccm.201212-2277OC (2013).

22. Jochems, S. P. et al. Inflammation induced by influenza virus impairs human innate immune control of pneumococcus. Nat Immunol 19, 1299–1308, doi:10.1038/s41590-018-0231-y (2018).

23. Wright, A. K. et al. Human nasal challenge with Streptococcus pneumoniae is immunising in the absence of carriage. PLoS Pathog 8, e1002622, doi:10.1371/journal.ppat.1002622 (2012).

24. Trevejo-Nunez, G., Elsegeiny, W., Conboy, P., Chen, K. & Kolls, J. K. Critical Role of IL-22/IL22-RA1 Signaling in Pneumococcal Pneumonia. J Immunol 197, 1877–1883, doi:10.4049/jimmunol.1600528 (2016).

25. Fais, S. et al. HLA-DR antigens on colonic epithelial cells in inflammatory bowel disease: I. Relation to the state of activation of lamina propria lymphocytes and to the epithelial expression of other surface markers. Clin Exp Immunol 68, 605–612 (1987).

26. Propst, S. M., Denson, R., Rothstein, E., Estell, K. & Schwiebert, L. M. Proinflammatory and Th2-derived cytokines modulate CD40-mediated expression of inflammatory mediators in airway epithelia: implications for the role of epithelial CD40 in airway inflammation. J Immunol 165, 2214–2221 (2000).

27. Cagnoni, F. et al. CD40 on adult human airway epithelial cells: expression and proinflammatory effects. J Immunol 172, 3205–3214 (2004).

28. Frick, A. G. et al. Haemophilus influenzae stimulates ICAM-1 expression on respiratory epithelial cells. J Immunol 164, 4185–4196 (2000).

29. Tettelin, H. et al. Complete genome sequence of a virulent isolate of Streptococcus pneumoniae. Science 293, 498–506, doi:10.1126/science.1061217 (2001).

30. Asmat, T. M., Agarwal, V., Rath, S., Hildebrandt, J. P. & Hammerschmidt, S. Streptococcus pneumoniae infection of host epithelial cells via polymeric immunoglobulin receptor transiently induces calcium release from intracellular stores. J Biol Chem 286, 17861–17869, doi:10.1074/jbc.M110.212225 (2011).

31. Peter, A. et al. Localization and pneumococcal alteration of junction proteins in the human alveolar-capillary compartment. Histochem Cell Biol 147, 707–719, doi:10.1007/s00418-017-1551-y (2017).

32. Herta, T. et al. DNA-release by Streptococcus pneumoniae autolysin LytA induced Krueppel-like factor 4 expression in macrophages. Sci Rep 8, 5723, doi:10.1038/s41598-018-24152-1 (2018).

33. Malley, R. et al. Recognition of pneumolysin by Toll-like receptor 4 confers resistance to pneumococcal infection. Proc Natl Acad Sci U S A 100, 1966–1971, doi:10.1073/pnas.0435928100 (2003).

34. Takahashi, S. et al. Pneumococcal Infection Aggravates Elastase-Induced Emphysema via Matrix Metalloproteinase 12 Overexpression. J Infect Dis 213, 1018–1030, doi:10.1093/infdis/jiv527 (2016).

35. Pollara, G. et al. Validation of Immune Cell Modules in Multicellular Transcriptomic Data. PLoS One 12, e0169271, doi:10.1371/journal.pone.0169271 (2017).

36. Schmeck, B. et al. Streptococcus pneumoniae induced c-Jun-N-terminal kinase- and AP-1-dependent IL-8 release by lung epithelial BEAS-2B cells. Respir Res 7, 98, doi:10.1186/1465-9921-7-98 (2006).

37. Siegel, S. J. & Weiser, J. N. Mechanisms of Bacterial Colonization of the Respiratory Tract. Annu Rev Microbiol 69, 425–444, doi:10.1146/annurev-micro-091014-104209 (2015).

38. Mahdi, L. K., Ogunniyi, A. D., LeMessurier, K. S. & Paton, J. C. Pneumococcal virulence gene expression and host cytokine profiles during pathogenesis of invasive disease. Infect Immun 76, 646–657, doi:10.1128/IAI.01161-07 (2008).

39. Briles, D. E., Novak, L., Hotomi, M., van Ginkel, F. W. & King, J. Nasal colonization with Streptococcus pneumoniae includes subpopulations of surface and invasive pneumococci. Infect Immun 73, 6945–6951, doi:10.1128/IAI.73.10.6945-6951.2005 (2005).

40. Morpeth, S. C. et al. Detection of Pneumococcal DNA in Blood by Polymerase Chain Reaction for Diagnosing Pneumococcal Pneumonia in Young Children From Low- and Middle-Income Countries. Clin Infect Dis 64, S347–S356, doi:10.1093/cid/cix145 (2017).

41. Talbot, U. M., Paton, A. W. & Paton, J. C. Uptake of Streptococcus pneumoniae by respiratory epithelial cells. Infect Immun 64, 3772–3777 (1996).

42. Jones, B. D., Paterson, H. F., Hall, A. & Falkow, S. Salmonella typhimurium induces membrane ruffling by a growth factor-receptor-independent mechanism. Proc Natl Acad Sci U S A 90, 10390–10394 (1993).

43. Gewirtz, A. T., Navas, T. A., Lyons, S., Godowski, P. J. & Madara, J. L. Cutting edge: bacterial flagellin activates basolaterally expressed TLR5 to induce epithelial proinflammatory gene expression. J Immunol 167, 1882–1885 (2001).

44. Hornef, M. W., Wick, M. J., Rhen, M. & Normark, S. Bacterial strategies for overcoming host innate and adaptive immune responses. Nat Immunol 3, 1033–1040, doi:10.1038/ni1102-1033 (2002).

45. Koppe, U. et al. Streptococcus pneumoniae stimulates a STING- and IFN regulatory factor 3-dependent type I IFN production in macrophages, which regulates RANTES production in macrophages, cocultured alveolar epithelial cells, and mouse lungs. J Immunol 188, 811–817, doi:10.4049/jimmunol.1004143 (2012).

46. Wilson, R. et al. Protection against Streptococcus pneumoniae lung infection after nasopharyngeal colonization requires both humoral and cellular immune responses. Mucosal Immunol 8, 627–639, doi:10.1038/mi.2014.95 (2015).

47. Zafar, M. A., Kono, M., Wang, Y., Zangari, T. & Weiser, J. N. Infant Mouse Model for the Study of Shedding and Transmission during Streptococcus pneumoniae Monoinfection. Infect Immun 84, 2714–2722, doi:10.1128/IAI.00416-16 (2016).

48. Plotkowski, M. C., Puchelle, E., Beck, G., Jacquot, J. & Hannoun, C. Adherence of type I Streptococcus pneumoniae to tracheal epithelium of mice infected with influenza A/PR8 virus. Am Rev Respir Dis 134, 1040–1044, doi:10.1164/arrd.1986.134.5.1040 (1986).

49. Pittet, L. A., Hall-Stoodley, L., Rutkowski, M. R. & Harmsen, A. G. Influenza virus infection decreases tracheal mucociliary velocity and clearance of Streptococcus pneumoniae. Am J Respir Cell Mol Biol 42, 450–460, doi:10.1165/rcmb.2007-0417OC (2010).

50. Zahlten, J. et al. Role of Pneumococcal Autolysin for KLF4 Expression and Chemokine Secretion in Lung Epithelium. Am J Respir Cell Mol Biol 53, 544–554, doi:10.1165/rcmb.2014-0024OC (2015).

51. Zahlten, J. et al. TLR9- and Src-dependent expression of Krueppel-like factor 4 controls interleukin-10 expression in pneumonia. Eur Respir J 41, 384–391, doi:10.1183/09031936.00196311 (2013).

52. Hotomi, M., Yuasa, J., Briles, D. E. & Yamanaka, N. Pneumolysin plays a key role at the initial step of establishing pneumococcal nasal colonization. Folia Microbiol (Praha) 61, 375–383, doi:10.1007/s12223-016-0445-z (2016).

53. Martner, A., Dahlgren, C., Paton, J. C. & Wold, A. E. Pneumolysin released during Streptococcus pneumoniae autolysis is a potent activator of intracellular oxygen radical production in neutrophils. Infect Immun 76, 4079–4087, doi:10.1128/IAI.01747-07 (2008).

54. Parker, D. et al. Streptococcus pneumoniae DNA initiates type I interferon signaling in the respiratory tract. MBio 2, e00016–00011, doi:10.1128/mBio.00016-11 (2011).

55. Bootsma, H. J., Egmont-Petersen, M. & Hermans, P. W. Analysis of the in vitro transcriptional response of human pharyngeal epithelial cells to adherent Streptococcus pneumoniae: evidence for a distinct response to encapsulated strains. Infect Immun 75, 5489–5499, doi:10.1128/IAI.01823-06 (2007).

56. Novick, S. et al. Adhesion and invasion of Streptococcus pneumoniae to primary and secondary respiratory epithelial cells. Mol Med Rep 15, 65–74, doi:10.3892/mmr.2016.5996 (2017).

57. Browall, S. et al. Intraclonal variations among Streptococcus pneumoniae isolates influence the likelihood of invasive disease in children. J Infect Dis 209, 377–388, doi:10.1093/infdis/jit481 (2014).

58. McCool, T. L., Cate, T. R., Moy, G. & Weiser, J. N. The immune response to pneumococcal proteins during experimental human carriage. J Exp Med 195, 359–365 (2002).

59. Hyams, C., Camberlein, E., Cohen, J. M., Bax, K. & Brown, J. S. The Streptococcus pneumoniae capsule inhibits complement activity and neutrophil phagocytosis by multiple mechanisms. Infect Immun 78, 704–715, doi:10.1128/IAI.00881-09 (2010).

60. Collins, A. M. et al. First human challenge testing of a pneumococcal vaccine. Double-blind randomized controlled trial. Am J Respir Crit Care Med 192, 853–858, doi:10.1164/rccm.201503-0542OC (2015).

61. Jochems, S. P. et al. Novel Analysis of Immune Cells from Nasal Microbiopsy Demonstrates Reliable, Reproducible Data for Immune Populations, and Superior Cytokine Detection Compared to Nasal Wash. PLoS One 12, e0169805, doi:10.1371/journal.pone.0169805 (2017).

62. Bray, N. L., Pimentel, H., Melsted, P. & Pachter, L. Near-optimal probabilistic RNA-seq quantification. Nat Biotechnol 34, 525–527, doi:10.1038/nbt.3519 (2016).

63. Soneson, C., Love, M. I. & Robinson, M. D. Differential analyses for RNA-seq: transcript-level estimates improve gene-level inferences. F1000Res 4, 1521, doi:10.12688/f1000research.7563.2 (2015).

64. Varet, H., Brillet-Gueguen, L., Coppee, J. Y. & Dillies, M. A. SARTools: A DESeq2- and EdgeR-Based R Pipeline for Comprehensive Differential Analysis of RNA-Seq Data. PLoS One 11, e0157022, doi:10.1371/journal.pone.0157022 (2016).

65. Chakraborty, S., Datta, S. & Datta, S. Surrogate variable analysis using partial least squares (SVA-PLS) in gene expression studies. Bioinformatics 28, 799–806, doi:10.1093/bioinformatics/bts022 (2012).

66. Fang, H., Knezevic, B., Burnham, K. L. & Knight, J. C. XGR software for enhanced interpretation of genomic summary data, illustrated by application to immunological traits. Genome Med 8, 129, doi:10.1186/s13073-016-0384-y (2016).

67. Kwon, A. T., Arenillas, D. J., Worsley Hunt, R. & Wasserman, W. W. oPOSSUM-3: advanced analysis of regulatory motif over-representation across genes or ChIP-Seq datasets. G3 (Bethesda) 2, 987–1002, doi:10.1534/g3.112.003202 (2012).

