## Supplemental Information for "Epithelial micro-invasion by *Streptococcus pneumoniae* induces epithelial-derived innate immunity during colonisation at the human mucosal surface"

#### SUPPLEMENTARY MATERIAL

| Antibody, Cat No. | Company, clone | Antibody | µl per sample | Biolegend Isotype | Filter on LSRII | Voltage run on LSRII |
|---|---|---|---|---|---|---|
| TCRγδ – PE-CF594 | BD, B1 | Mouse/IgG1,k | 5 | n/a | 610/20 | 500 |
| IL22Ra1-PerCP | R&D | Mouse/IgG1 | 5 | #400147 | 695/40 | 661 |
| CD54 –BV711 | MAH-HA58, BD | Mouse/IgG1,k | 5 | #400167 | 710/50 | 615 |
| CD107a –BV650 | Biolegend, H4A3 | Mouse/IgG1,k | 5 | n/a | 655/8 | 492 |
| CD3-APC.Cy7 | BD, SK7 | Mouse/IgG1,k | 3 | n/a | 780/60 | 710 |
| CD45-PacOrange | Life technologies, HI30 | Mouse / IgG1 | 3 | n/a | 525/50 | 532 |
| CD4 –BV605 | Biolegend, RPA-T4 | Mouse IgG1,k | 3 | n/a | 610/20 | 500 |
| CD40-PE.Cy7 | Biolegend, 5C3 | Mouse/IgG1,k | 3 | #400125 | 780/60 | 693 |
| CD218a-APC | eBiosciences, H44 | Mouse/IgG1,k | 5 | n/a | 660/20 | 542 |
| EpCAM-PE | Biolegend, 9C4 | Mouse/IgG2b,k | 1.5 | #401207 | 575/26 | 488 |
| CD8 –BV785 | Biolegend SK1 | Mouse/IgG1,k | 1.5 | n/a | 780/60 | 591 |
| HLA-DR-FITC | Biolegend L243 | Mouse/ IgG2a,k | 0.75 | #400207 | 530/30 | 400 or 500 |
| IL-22RA1-PerCP | Bio Techne Ltd, 305405 | Mouse/IgG1 | 5 | #400147 | 695/40 | 551 |
| IL-22RA1-PerCP | Bio Techne Ltd, 305405 | Mouse/IgG1 | 5 | #400147 | 710/50 | 551 |
| CD54 –APC | Life Technologies 1H4 | Mouse/IgG1,k | 5 | #555745 | 670/30 | 592 |
| CD107a –PE | Life Technologies H4A3 | Mouse/IgG1,k | 5 | #555749 | 582/15 | 510 |
| CD40-BV421 | Biolegend, 5C3 | Mouse/IgG1,k | 5 | #562438 | 450/50 | 528 |
| HLA-DR-BUV395 | BD, G46-6 | Mouse/ IgG2a,k | 5 | #563809 | 379/28 | 538 |

**Panel composition and specifications of the multiparametric flow cytometry panel used for the identification of the epithelial and immune cells from nasal cells, and for the epithelial activation markers from Detroit 562 cell lines**. A multiparametric flow cytometry panel composed of twelve (*in vivo*) and five (*in vitro,* last five) different monoclonal antibodies was developed to analyze the epithelial activation and immunological cell populations obtained from nasal cell samples by flow cytometry. Electronic compensation was set using CompBeads (BD Biosciences) and the FacsDiva automated compensation according to manufacturer's instructions.

| Antibody | Company, clone |
|---|---|
| JAM-A | Santa Cruz, 1H2A9 |
| Claudin 4 | Life Technologies |
| β catenin | NEBiolabs Ltd, L54E2 |
| Pneumococcal Antiserum Type 4 | SSI Diagnostica (TIGR4) |
| Pneumococcal Antiserum Pool Q | SSI Diagnostica (6B and 23F) |
| Wheat Germ Agglutinin | Vector labs, Rhodamine conjugated |
| Goat anti-mouse | Life Technologies (AF- 546,647) |
| Goat anti rabbit | Life Technologies (AF-488.546,647) |
| Goat anti-mouse-HRP | Dako (P0047) |

**Panel of Antibodies for Immunofluorescence.** Samples were incubated with primary antibodies for one hour (1:100). After washing, samples were incubated with secondary antibodies (1 : 500) for 45 minutes.

#### *Ex vivo* cell culture

Primary epithelial cells were cultured according to Müller et al., Jove, 2013 with modification[1]. Briefly, nasal cell curette biopsy samples were collected in RPMI containing Penicillin-Streptomycin (1%) and FBS (10%) before centrifugation at 400g for 5 minutes. Cells were resuspended in BEGM-AS [25ml PneumaCult BEGM (Bronchial Epithelial Growth Media, Stem Cell Technologies), 25ml BEGM (Cell Application), GA-1000 (1:2000, Lonza)] supplemented with 50ng/ml DNase I), incubated for 20min at RT and centrifuged. The resuspended pellet containing BEGM-$S^{++}$ media [50ml PneumaCult BEGM media, Sodium Bicarbonate (1:50, Fisher Scientific) Nu-serum IV (1:20), GA-1000 (1:1000)], was plated into 12 well cell culture plates (Corning) that had been coated with 300µg/ml PureCol

(Cell System) for 30minutes at 37°C and washed with Hanks Buffer Saline Solution ($HBSS^{+/+}$, with calcium and magnesium, Gibco). 24 hours after sampling, media was replaced with BEGM-$S^{++}$ and BEGM-$AS^{+}$ [50ml of BEGM-AS media, 2nM retinoic acid, GA-1000 (1:2000)] and applied as follows: 24h - 1:1; 48h - 1:3, 72h and following - 1:6. At day 7, the cells were dislodged using 0.25% trypsin, placed in a prepared 15ml flacon containing Soy trypsin inhibitor (Sigma Aldrich), centrifuged (4°C, 400g for 5 minutes) and resuspended in BEGM-$AS^{+}$ and seeded into a cell culture flask (Corning). Cells were cultured for a maximum of 20 days. At day 28 (Week 4) cells were seeded on Corning 12 well inserts (culture area = 1.12$cm^2$) and cultured in BEGM ALI media (25 ml DMEM high glucose, 25 ml BEGM-AS media, 50 µg/ml Bovine Pituitary Extract, 66 µg/ml Nystatin (Fisher Scientific), 1.5 µg/ml BSA, GA-1000 (1:4000)). Cells were seeded at a concentration of $1x10^5$ cells/well and cultured until confluent. Apical and basolateral media was replaced with BEGM ALI media supplemented with 500nM retinoic acid for 48 hrs. Media was then replaced with PneumaCult-ALI media on the basal side only. Cells were maintained for 30 days with the apical cell surface rinsed weekly with 10 minutes of $HBSS^{+/+}$ and removing liquid thereafter.

**Electron Microscopy**

Transmission: Cells were fixed in 2% paraformaldehyde and 1.5% glutaraldehyde in 0.1M sodium cacodylate buffer pH7.3 and incubated for 24 hours at $3^oC$. Samples are washed twice in 0.1M sodium cacodylate for 30 minutes and post-fixed in 1% Oso4 / 1.5% K4Fe(CN)6 in 0.1M cacodylate buffer pH7.3 for 1 hour at $3^oC$. Samples are washed in distilled water for 5 minutes. The cells were dehydrated in a graded ethanol-water series and infiltrated with Agar 100 epoxy resin by the following steps: 25 % Ethanol 5 minutes; 50 % Ethanol 5 minutes; 70% Ethanol 5 minutes; 90% Ethanol 5 minutes; 100 % ethanol (dry) 5 minutes, four times. This was followed by incubation in propylene oxide for 5 minutes; 2:1

propylene oxide: resin for one hour; 1:1 propylene oxide 1 parts resin mix x 1 hr; 1:2 propylene oxide: resin for one hour; 100 % resin for 4 hours; 100% resin for 16 hours. The Epoxy Resin was a medium hardness mix and the Agar resin consisted of Agar 100 (12 gm), DDSA (8 gm), MNA (5 gm) and BDMA (0.65 ml).

The samples were hardened ready to section by placing the coverslip cell-side down onto a resin-filled beam capsule and hardened at 60$^{o}$c for 48 hours. The coverslip was removed and a representative area was selected. Ultra-thin sections were cut at 70-80 nm using a diamond knife on a Reichert ultra-cut S microtome. Sections were collected on copper grids and stained with lead citrate. Samples were viewed with a Joel 1010 transition electron microscope and the images recorded using a Gatan Orius CCD camera.

Scanning: The samples were post-fixed in 1% Oso4/ 1.5% K4Fe (CN)6 in 0.1M cacodylate buffer at 3$^{o}$C for 1.5 hours. Samples were washed in 0.1M cacodylate buffer, rinsed in distilled water and dehydrated in a graded ethanol-water series to 100% ethanol. (25 % Ethanol 5 minutes; 50 % Ethanol 5 minutes; 70% Ethanol 5 minutes; 90% Ethanol 5 minutes and finally 100 % ethanol (Annular and dry) 5 minutes, four times). Samples were critical point dried using $CO_2$ and mounted on aluminium stubs using sticky carbon taps. The mounted samples were coated with a thin layer of ~10nm thick Au. using a sputter coater and viewed and images recorded with a FEI.

#### SUPPLEMENTARY FIGURE LEGENDS

**Supplementary Excel File 1.** Differentially upregulated genes from Detroit 562 cells infected with pneumococci; comparisons against non-infected cells.

**Supplementary Excel File 2.** REACTOME Pathways and Interactome Genes from Detroit 562 cells infected with pneumococci.

**Supplementary Excel File 3.** Transcription factor binding site enrichment analysis

**Supplementary Excel File 4.** Differential gene expression from the EHPC model

**Supplementary Figure 1. *Streptococcus pneumoniae* colonisation of the human nasal mucosa is associated with adhesion, micro-colony formation and microinvasion**

(a) *In vivo* Immunofluorescence: Cells collected from nasal curette biopsies from the EHPC model were prepared for microscopy and stained for surface carbohydrates (red), 6B (green) and DAPI (blue). In addition to epithelial cells collected from nasal biopsies, immune cells were readily observed as indicated by round, multi-nuclei cells. (b) *In vivo* electron microscopy: cells collected from nasal curette biopsies from a natural carrier of the pneumococcus were prepared for TEM. MV – microvilli, N – nucleus, TJ – tight junctions, G – goblet cell, *S.pn* – *Streptococcus pneumoniae.* (i) epithelial cell architecture preserved, scale bar 2μm; (ii) intact sheet of cells with tight junctions visible, scale bar 2μm; (iii) a diplococci adhered to the cell surface, scale bar 2μm (insert 200nm); (iv) a pneumococcal chain, scale bar 0.5μm. (c) *Ex vivo* Immunofluorescence: primary cells were cultured for 30 days on an air-liquid interface and incubated with 1 million pneumococci (6B (i) or 23F (iii)) for 3 hours. (i) Cells were fixed and stained for β catenin (blue), JAM-A (red) and pneumococci capsule (green). Nuclei in blue. The XZ image is shown in (ii and iv, respectively). Images represent (v) intracellular, (vi) paracellular and (vii) basal localisation of *S. pneumoniae* within the epithelial monolayers. Images represent one experiment from one transwell insert per condition.

**Supplementary Figure 2. Epithelial surface marker expression in response to *Streptococcus pneumoniae in vivo***

(a-d) *In vivo* FACS: Cells collected from nasal scrapes were prepared for Flow Cytometry Analysis. (a) Compensation matrix for flow cytometry panel. (b) Sample sizes for each day and carriage status. Sample sizes refer to: (CD40, CD54, IL-22Ra1) (CD107a) (HLADR) respectively. (c) Samples were gated into 'all cells', 'single cells', and finally 'EpCAM positive' cells, against an empty channel (AF-700-A) to analyse the epithelial cell population of the samples. Representative plots are shown. (d) An example of histograms for each epithelial activation marker, from a carriage positive and carriage negative volunteer. From these histograms, the cells expressing the highest 5% of activation markers, from baseline, were gated (high), to generate comparisons between sampling days within volunteers. Median fluorescence intensity and high surface marker-expressing cells (≥95% of the baseline expression) for epithelial surface expression of (e) IL-22Ra, (f) HLADR and (g) CD40.  Results are from a minimum of two volunteers. Black circles show carriage negative, and grey circles show carriage positive, volunteers.

**Supplementary Figure 3. Junction association of *S. pneumoniae* in Detroit 562 cells**

(a) Detroit 562 cells were stained for *S. pneumoniae* and either WGA, JAM-A, Claudin 4 or β catenin three hours post infection. Representative XY images and XZ stacks of cells are shown for examples of surface 6B; basal located 23F; Micro-colony of TIGR4, surface bound dPly. N = 3 with replicates. (b) Negative controls for immunofluorescence (i) JAM-A (ms) primary antibody and goat-anti rabbit secondary with TIGR4-FAMSE. (ii) TIGR4-FAMSE and goat-anti mouse secondary. (iii) antiserum and goat anti-mouse secondary. (iv) blank insert with FAMSE-23F with goat anti-mouse and goat anti-rabbit secondary. (v) Blank insert with JAM-A and goat anti-mouse secondary. (vi) FAMSE-TIGR4 with JAM-A antibody.

**Supplementary Figure 4. Epithelial adherence, endocytosis and transmigration by *Streptococcus pneumoniae* varies by pneumococcal strain and is modulated by pneumolysin, without affecting barrier integrity.**

(a - f) A549 cells were stimulated with *S. pneumoniae* for three hours and CFU measured for (a) association, (b) internalisation and (c) transmigration over time. ****P = <0.0001 for adhesion and invasion assays; **P = 0.0049 at three hours transmigration for 23F v TIGR4 and TIGR4 v dPLY. N = 7. (d) Cells stained for *S. pneumoniae* (green) and either WGA, JAM-A, Claudin 4 or β catenin (red). XY and XZ images represent associations and destinations of the different pneumococcal strains with different host proteins. (e – f) Barrier function was assessed before and after three hours exposure to *S. pneumoniae.* (e) There were no significant differences in TEER at the start of the experiment. N = 4. ****P = <0.0001. (f) Dextran leakage was recorded. Blank inserts = positive control (****P < 0.0001). *P = 0.0108 non-infected v TIGR4. n = 5.

(g - l) Calu3 cells were stimulated with *S. pneumoniae* for three hours and CFU measured for (g) association, (h) internalisation and (i) transmigration over time. ****P = <0.0001 for adhesion and invasion. (i) Transmigration at three hours infection between 23F v TIGR4 *P = 0.0281. N = 4. (j) Calu3 cells stained for *S. pneumoniae* and either WGA, JAM-A, Claudin 4 or β catenin. XY and XZ images represent associations and destinations of the different pneumococcal strains with different host proteins. (k - l) Barrier function was assessed during exposure to *S. pneumoniae.* (k) TransEpithelial Electrical Resistance (TEER) was recorded before and after the experiment: n = 7. TEER was not significantly different between inserts before or after pneumococci were added (P = 0.7507 and 0.1088 respectively). (l) Dextran leakage was measured and calcium withdrawal = positive control

(****P = <0.0001, n = 3, Unpaired T-Test). No significant differences were detected in permeability with pneumococcal infection, P = 0.0560. N = 5.

**Supplementary Figure 5. Internalised pneumococci egress from the epithelium**

(a) Detroit 562 cells were incubated with *S. pneumoniae* for three hours, washed, treated with gentamicin for 1 hour, washed and the cultures incubated for further time points to measure (a) bacterial internalisation. N = 4. (b) Bacteria that were released into the apical or basal chamber after one hour were then counted. N = 3. Similar results were also observed with Calu 3 cells (data not shown).

**Supplementary Figure 6. Epithelial surface marker expression in response to *Streptococcus pneumoniae in vitro***

(a - c) *In vitro* FACS: Cells collected from Detroit 562 monolayers (compensation matrix shown in (a)) were gated into ‘all cells’ (b), the population further defined to ‘single cells’, and finally exclusion of dead cells (defined by treatment with $H_2O_2$, data not shown), lead to the live cell population used for analyses. (c) Representative histograms for each epithelial activation marker are shown. Median fluorescence intensity and cells expressing the highest 5% of surface expression for (d) IL-22Ra, (e) HLADR and (f) CD40 were gated to generate comparisons between strains of pneumococci and non-infected cells.

**Supplementary Figure 7. Pneumolysin activity is comparable between pneumococcal strains.**

(a - c) Bacterial preparations in phenol free RPMI (Invitrogen) were added to a solution of 2% red blood cells (EO labs) in a U-bottom 96 well plate for 30 minutes at 37$^{o}$C / 5% $CO_2$. There were no significant differences between pneumococcal strain density for each dilution.

(b) Serial dilutions of 0.5% saponin was used as a positive control for cell lysis. The plate was centrifuged for 1 minute at 1000g to pellet the un-lysed red blood cells and supernatant absorbance was read at 540nm. (c) There were no differences in lysis potential between the pneumococcal strains. dPly-TIGR4 mutant was used as a negative control for lysis.

**Supplementary Figure 8. Epithelial transcriptomic responses to *Streptococcus pneumoniae in vitro***

Individual pathways represented in the clusters from Figure 6c.

#### REFERENCES

1. Muller, L., Brighton, L.E., Carson, J.L., Fischer, W.A., 2nd & Jaspers, I. Culturing of human nasal epithelial cells at the air liquid interface. *J Vis Exp* (2013).

#### Supplementary Figure 1

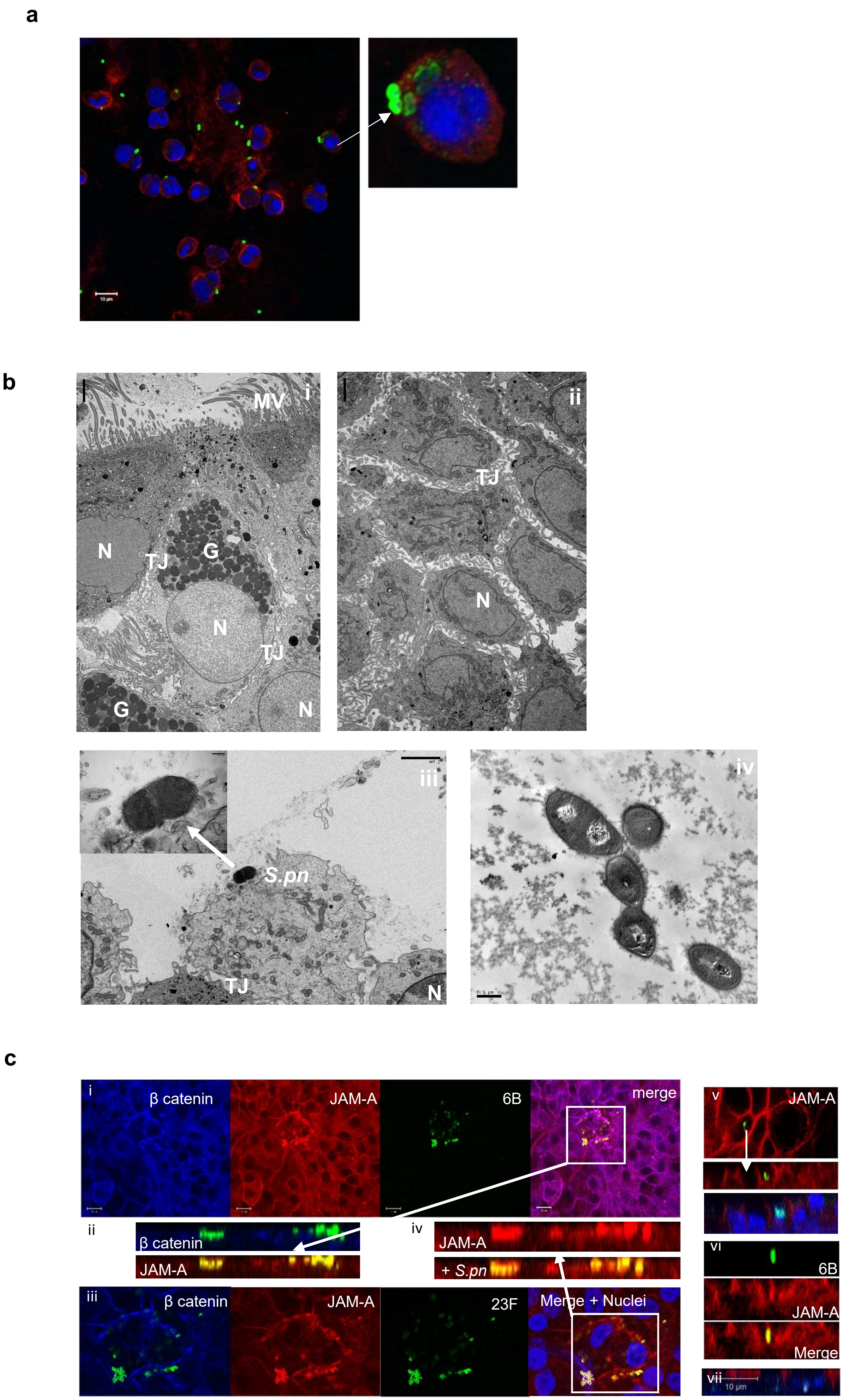

### Supplementary Figure 2

**a**

[M] Edit Save Matrix SSM

| Show All | FITC-A :: HLAD... | PE-A :: Epcam, ... | PE-Texas Red-... | PerCP-Cy5-5-A... | PE-Cy7-A :: CD... | Pacific Blue-A :: ... | Pacific Orange-... | BV605-A :: CD4 | BV650-A :: CD1... | BV711-A :: CD5... | BV786-A :: CD8 | APC-A :: CD218... | Alexa Fluor 700-A | APC-Cy7-A :: C... |
|---|---|---|---|---|---|---|---|---|---|---|---|---|---|---|
| FITC-A :: HLADR, CD66b | 100 | 13.8158 | 6.2607 | 1.2509 | 0.1647 | 0.0115 | 4.5169 | 0.2008 | 0.0386 | 0.0232 | 0.0057 | 0.0077 | 0 | 0.0113 |
| PE-A :: Epcam, EpCam | 1.5588 | 100 | 41.8032 | 10.9439 | 1.3986 | 0 | 0.1784 | 2.568 | 0.5518 | 0.2166 | 0.0608 | 0.0068 | 0 | 0 |
| PE-Texas Red-A :: Tgd, TCRgd, ... | 0.258 | 8.7632 | 100 | 39.5663 | 5.9608 | 0.0078 | 0.0486 | 6.2893 | 1.4458 | 0.7501 | 0.3323 | 0.2284 | 0.07 | 0.0725 |
| PerCP-Cy5-5-A :: IL22RA, IL22Ra... | 0 | 0.0132 | 0.0151 | 100 | 15 | 0 | 0.0364 | 0.0099 | 1 | 7 | 4.8525 | 3.5 | 5.7058 | 4.5119 |
| PE-Cy7-A :: CD40, HLA-DR | 0.1204 | 0.6791 | 0.3295 | 0.2717 | 100 | 0 | 0.0193 | 0.0211 | 1 | 0.0088 | 3.1261 | 1 | 0.3884 | 6.7784 |
| Pacific Blue-A :: Viability, Live D... | 0.6054 | 0.0625 | 0.0292 | 0 | 0.017 | 100 | 153.2052 | 3.185 | 0.6535 | 0.2589 | 0.14 | 0.0785 | 0 | 0 |
| Pacific Orange-A :: CD45 | 0.8344 | 0.4321 | 0.3725 | 0.149 | 0 | 0.3069 | 100 | 25.8392 | 7.7307 | 3.4337 | 1.4771 | 0.0633 | 0 | 0.0067 |
| BV605-A :: CD4 | 0.0085 | 1.136 | 13.6955 | 6.3801 | 1.3388 | 1.0337 | 0.8962 | 100 | 36.7231 | 15.2 | 7.178 | 0.3013 | 0.0927 | 0.0419 |
| BV650-A :: CD107a, CD107 | 0.0836 | 0.0767 | 0.5133 | 4.4226 | 0.7271 | 1.9928 | 1.5505 | 15.7727 | 100 | 43 | 16.5365 | 31.6808 | 10.4587 | 5.4768 |
| BV711-A :: CD54, PD1 | 0.0485 | 0.0388 | 0.0194 | 34.9144 | 13 | 3.2091 | 2.616 | 0.1562 | 0.6688 | 100 | 77.1057 | 0.5443 | 43.6319 | 30.3307 |
| BV786-A :: CD8 | 0.0827 | 0.0489 | 0.0414 | 0.1091 | 13.5479 | 2.5174 | 3.0766 | 0.3293 | 0.2572 | 0.5812 | 100 | 0.0897 | 0.8839 | 10.8286 |
| APC-A :: CD218a, CD16 | 0.0083 | 0.0058 | 0.0219 | 1.3207 | 0.2895 | 0.0152 | 0.0417 | 0.0381 | 3.7194 | 1.25 | 0.734 | 100 | 30.8557 | 14.8498 |
| Alexa Fluor 700-A | 0 | 0 | 0 | 0 | 0 | 0 | 0 | 0 | 0 | 0 | 0 | 0 | 100 | 0 |
| APC-Cy7-A :: CD3 | 0.0352 | 0.0294 | 0.0176 | 0.0529 | 2.3721 | 0 | 0 | 0 | 0.0455 | 0.0303 | 3.6438 | 0.8585 | 3.4813 | 100 |

**b**

| | Baseline | Day 2 | Day 6 | Day 9 | Day 14-29 |
|---|---|---|---|---|---|
| Carriage negative | (5)(9)(14) | (3)(4)(11) | (3)(7)(17) | (3)(5)(15) | (1)(5)(15) |
| Carriage positive | (8)(10)(13) | (9)(13)(18) | (4)(9)(14) | (6)(11)(17) | (5)(8)(11) |

**c**

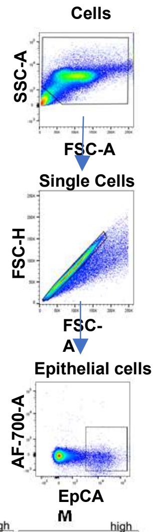


**d**

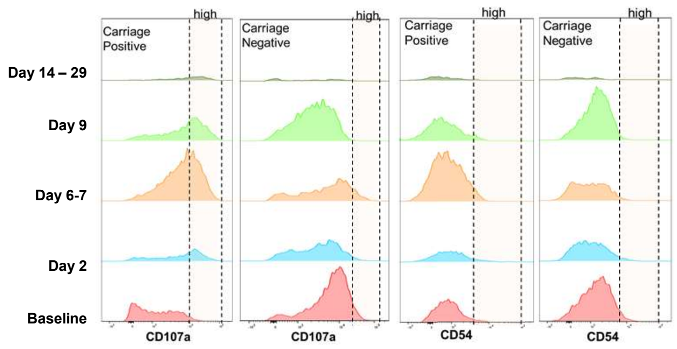
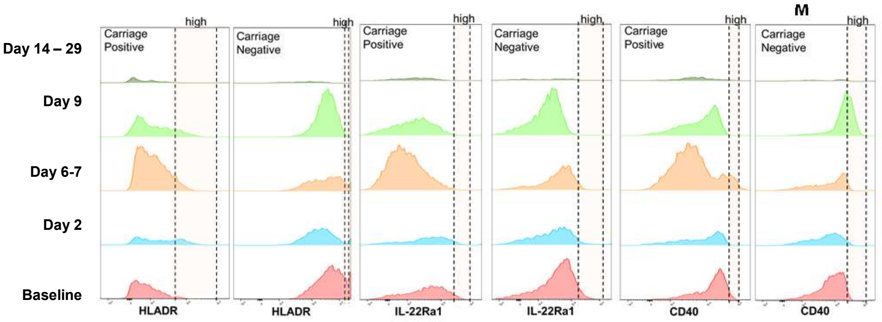


**e**

**f**

**g**

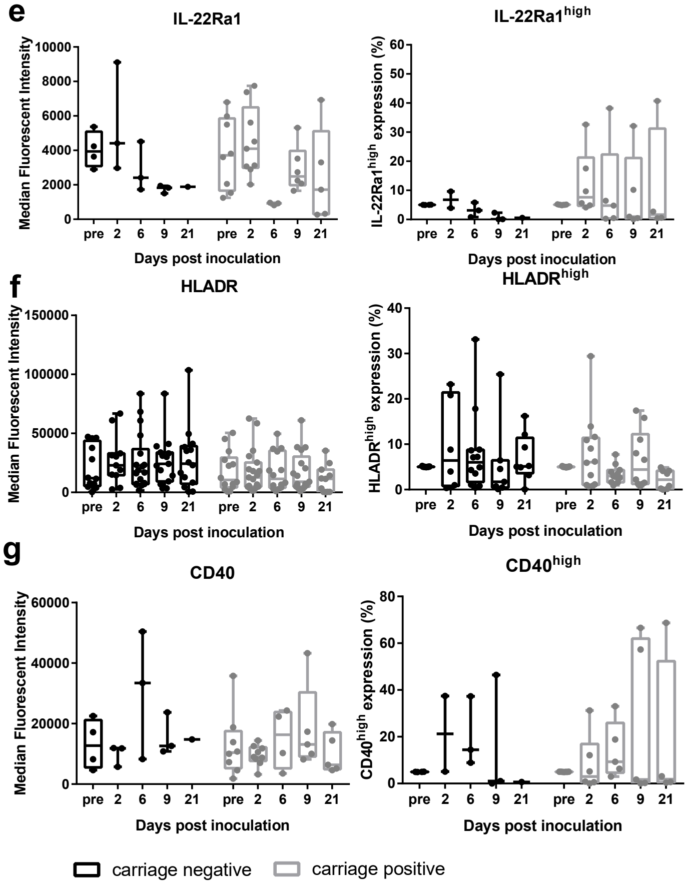

### Supplementary Figure 3

**a**

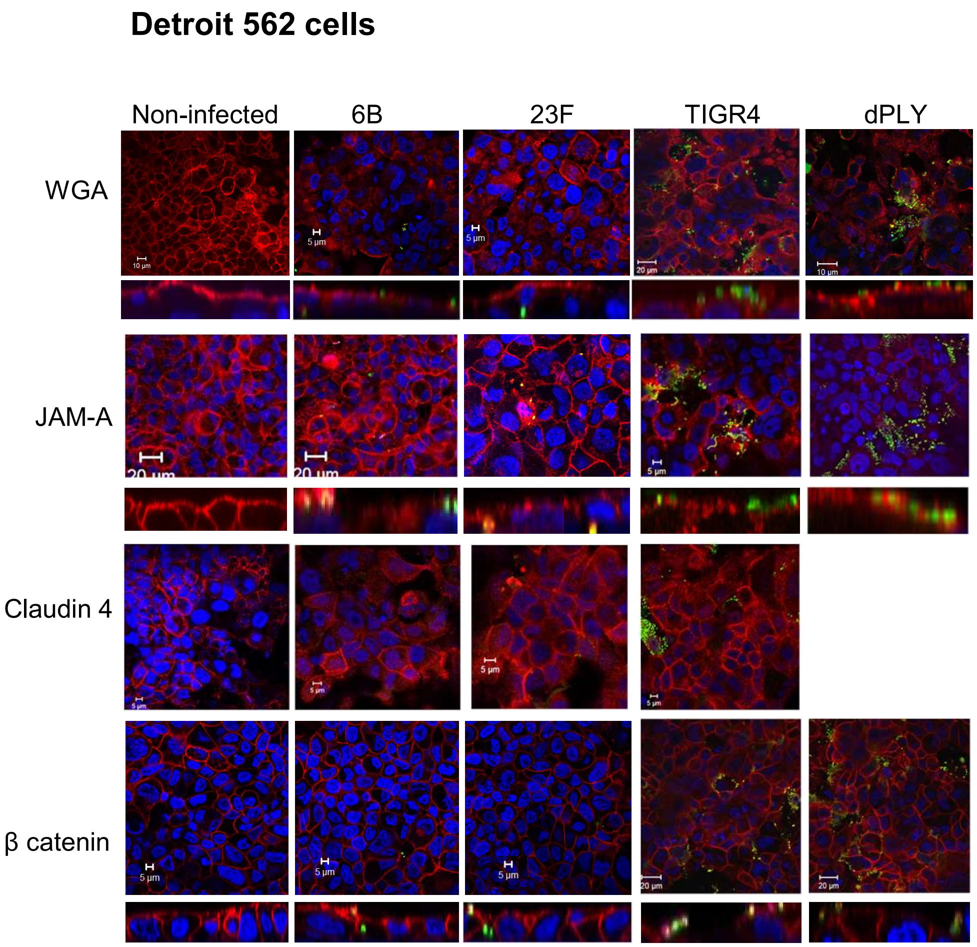


**b**

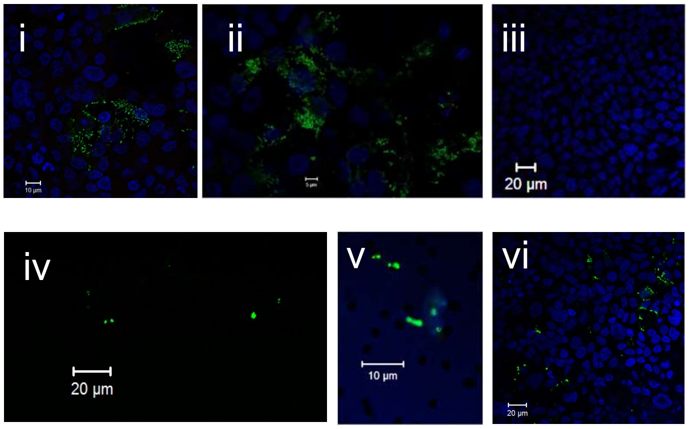

### Supplementary Figure 4 A549 cells

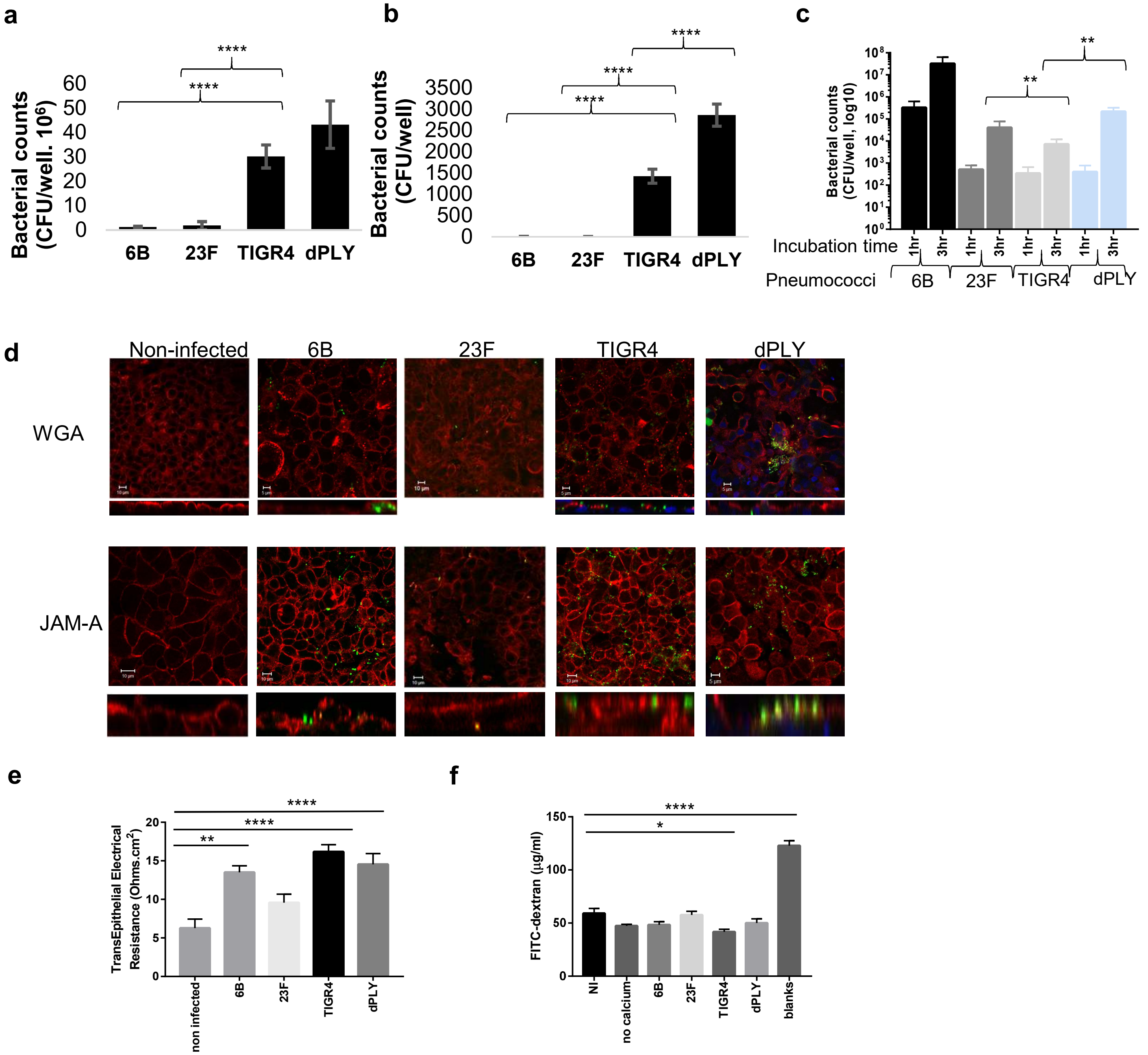

**Calu3 cells**

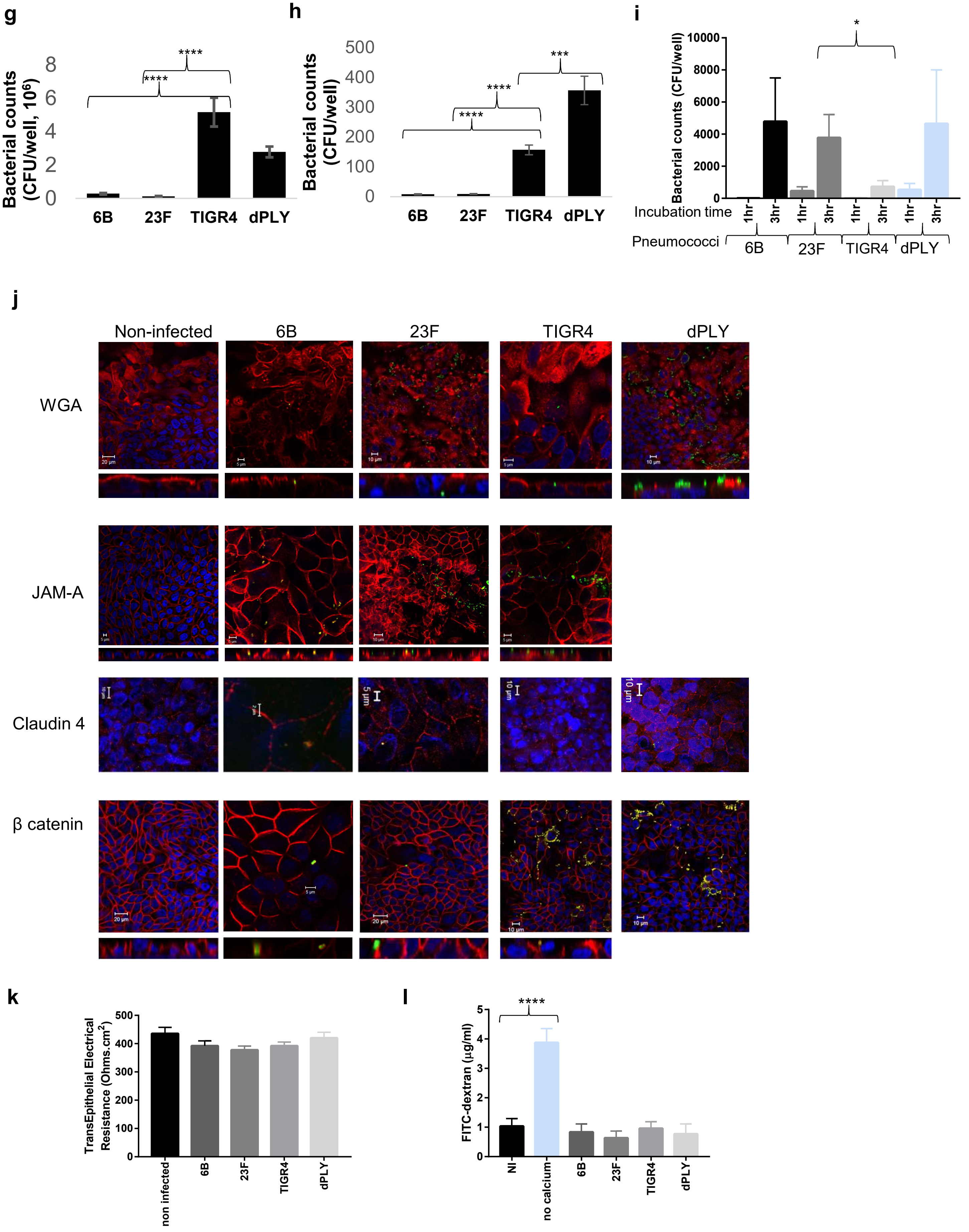
g
Bacterial counts (CFU/well, $10^6$)
0
2
4
6
8
****
****
6B
23F
TIGR4
dPLY
h
Bacterial counts (CFU/well)
0
100
200
300
400
500
****
****
***
6B
23F
TIGR4
dPLY
i
Bacterial counts (CFU/well)
0
2000
4000
6000
8000
10000
*
Incubation time
1hr
3hr
1hr
3hr
1hr
3hr
1hr
3hr
Pneumococci
6B
23F
TIGR4
dPLY
j
Non-infected
6B
23F
TIGR4
dPLY
WGA
JAM-A
Claudin 4
β catenin
k
TransEpithelial Electrical Resistance (Ohms.cm$^2$)
0
100
200
300
400
500
non infected
6B
23F
TIGR4
dPLY
l
FITC-dextran (µg/ml)
0
1
2
3
4
5
****
NI
no calcium
6B
23F
TIGR4
dPLY

### Supplementary Figure 5

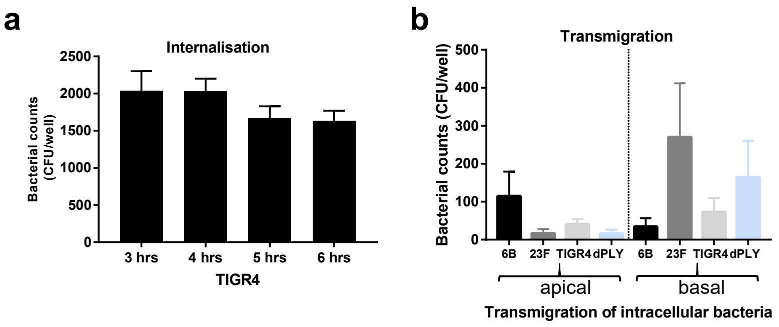

### Supplementary Figure 6

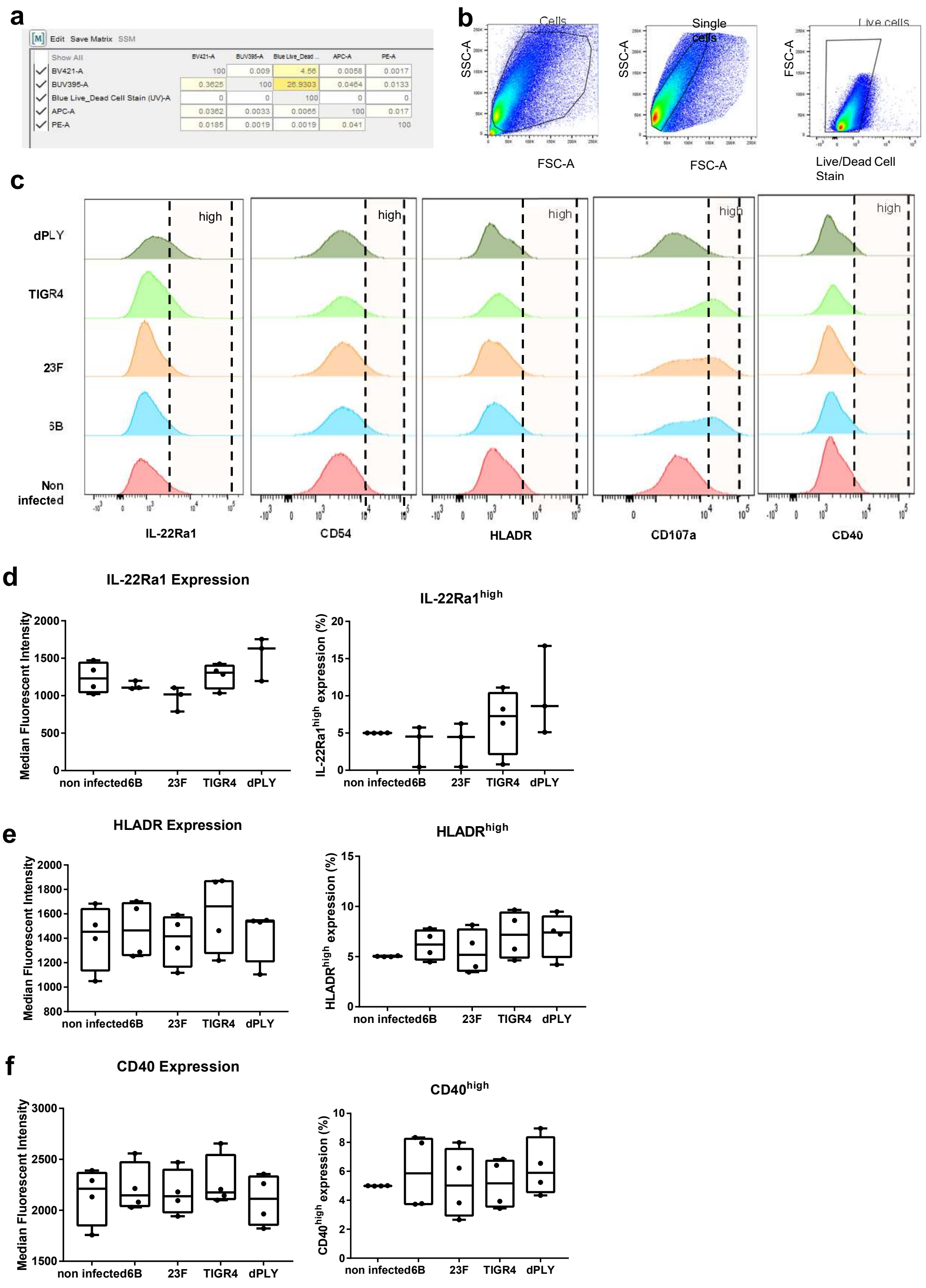

### Supplementary Figure 7

**a**

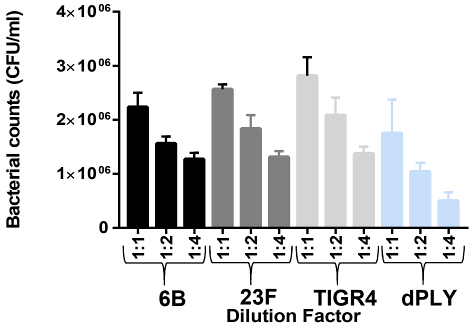


**b**

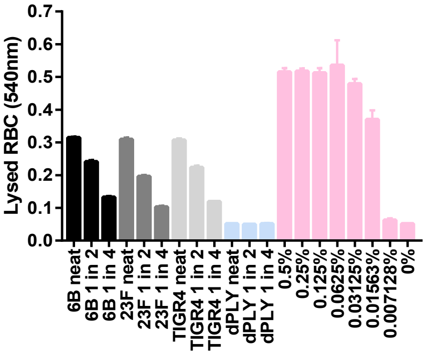

### Supplementary Figure 8

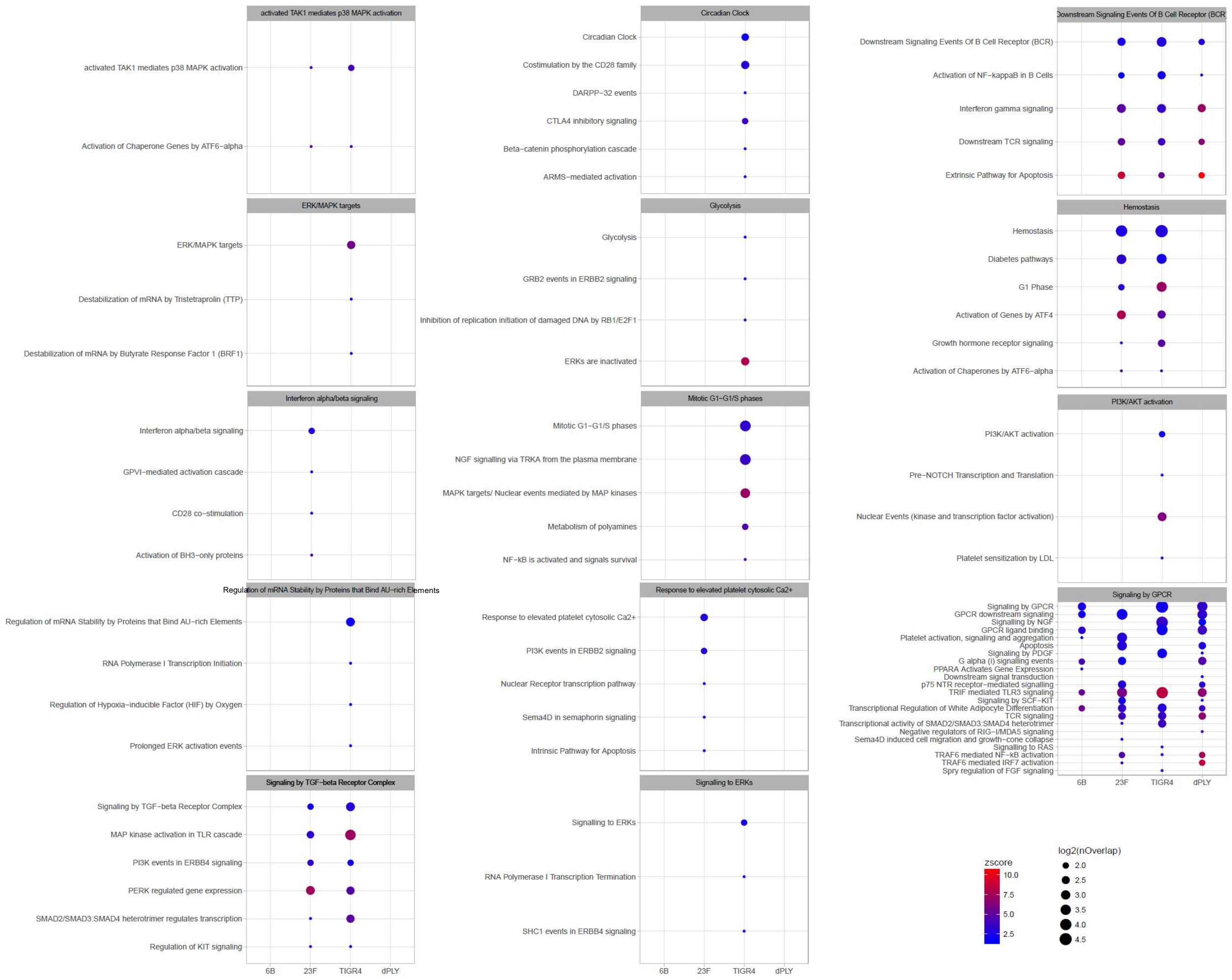
